# Paternal-effect genes revealed through semen cryopreservation in *Perca fluviatilis*

**DOI:** 10.1101/2023.12.06.570413

**Authors:** Abhipsa Panda, Sylwia Judycka, Katarzyna Palińska-Żarska, Rossella Debernardis, Sylwia Jarmołowicz, Jan Jastrzębski, Taina Rocha de Almeida, Maciej Błażejewski, Piotr Hliwa, Sławek Krejszeff, Daniel Żarski

## Abstract

Knowledge about paternal-effect genes (the expression of which in progeny is controlled by the paternal genome) in fish is very limited. To explore this issue, we used semen cryopreservation as a specific challenge test for sperm cells, thus enabling selection amidst cryo-sensitivity. We created two groups of Eurasian perch (*Perca fluviatilis*) as a model – eggs fertilized either with fresh (Fresh group) or cryopreserved (Cryo group) semen from the same male followed by zootechnical-transcriptomic examination of consequences of cryopreservation in obtained progeny (at larval stages). Most of the zootechnical observations were similar in both groups, except the final weight was higher in the Cryo group. Semen cryopreservation appeared to act as a “positive selection” factor, upregulating most paternal-effect genes in the Cryo group. Transcriptomics profile of freshly hatched larvae sourced genes involved in the development of visual perception as paternal-effect genes. Consequently, larvae from the Cryo group exhibited enhanced eyesight, potentially contributing to more efficient foraging and weight gain compared to the Fresh group. This study unveils, for the first time, the significant influence of the paternal genome on the development of the visual system in fish, highlighting *pde6g*, *opn1lw1*, and *rbp4l* as novel paternal-effect genes.

## Introduction

Fish ontogeny, especially at their early life stages, is largely determined by their parents. However, most of the work revolves around the evaluation of maternal contribution ^1^. This is mostly associated with the well-known fact that the mother provides nutritional and energy reserves (contained in the yolk) utilized by fish during the larval period ^2^. More recently, it became apparent that maternal contribution is far beyond the nutritional reserves, shedding light on the remaining components of egg molecular cargo (especially transcripts) as a modulator of progeny phenotype ^3^. Nevertheless, in the past three decades, a piling body of evidence indicates significant paternal influence during the early life history (ELH) (i.e., from fertilization until the end of the larval period in fish) ^4–7^. In addition to reproductive fitness and age of males, other wide variety of progeny traits, such as embryonic developmental rate and larval size upon hatching, have been shown to be contributed by the father ^8^. These finding sheds light on the fact that very little is known about the paternal contribution to progeny phenotype in fishes. Understanding paternal contribution to fish ELH is crucial for fish biologists, ecologists and the aquaculture sector, where it can be used to fine-tune selective breeding programs, which may lead to increased production effectiveness and improved welfare of cultured species ^9^.

Considering paternal effects in fishes, offspring characteristics are evidently not influenced solely by the haploid set of genes contributed by the father but also by the interaction between genes and the environment having, among others, a direct effect on epigenetic modifications ^5,10,11^. Extrinsic factors, however, very often affect particular genes selectively and do not affect the whole genome at the same level ^12^. In effect, some genes are susceptible to particular environmental cues, while others remain intact. Some portion of the epigenetically susceptible genes of the paternal genome may further affect the expression of this gene in developing embryos and consequently modify the progeny phenotype. This phenomenon of environment-dependent control over the phenotype will be hereinafter referred to as “nongenetic inheritance” ^13,14^. The genes whose expression in the progeny is controlled by the male gametes hereinafter will be referred to as “paternal-effect genes”. In layman language, one would understand paternal-effect genes as those genes that are present in both the oocyte’s and the spermatozoa’s genetic material, but the latter’s affect its expression in progeny.

While standardizing sophisticated reproductive techniques, semen cryopreservation (involving specific procedures enabling effective storage of the viable cells at ultra-low temperatures for a very long period) for fishes has come into limelight over several decades now ^15,16^. Cryobanking of semen helps to manage the genetic diversity of the fish species, facilitates spawning synchronization, allows selective breeding and much more ^17^. Cryopreservation of semen is a big shock to a sperm cell constituting a specific challenge test ^18^ which causes irreversible damages to the cryo-sensitive cells which lose fertilizing capacity, in contrast to cryo-resistant cells retaining their functionality. Importantly, the effect of cryopreservation on spermatozoa functionality depends on the species. On one hand, sperm motility, fertilization rate and the hatching rates were seen to be high and similar with post-thaw semen when compared with use of fresh semen in some Salmoniforms and Esociformes ^19–21^. On the other hand, studies done so far on few other teleosts (e.g., Levantine scraper, wild brown trout, Atlantic cod) report reduced motility and viability of sperm after freezing-thawing ^22^. Also, the post-thaw semen from the same fish when used for *in vitro* fertilization, there were significant declination in fertilization success when compared with fresh sperm control, followed by abnormalities like cleavage patterns, hatching success, organogenesis etc. ^23,24^. So far studies elaborating implications of sperm cryopreservation on embryonic and larval performance as well as gene expression profile in offspring is very limited ^18^. It has been reported, that along with DNA damage and changes in chromatin integrity in sperm, the transcriptome of the larvae (obtained with frozen-thawed sperm) is also seen to be compromised upon sperm cryopreservation ^25^. Even though after fertilization, the zygotic genome has the ability to repair a certain degree of the DNA damage, developmental failures can still be observed indicating that this process has obvious limitations, as seen in humans, cattle, fishes etc. ^26^. Clearly, semen cryopreservation seems to be a very subjective kind of sperm cell stressor that depends on the cell/species type.

It has recently been shown that the phenotype of the progeny is affected by the epigenetic profile inherited directly from the paternal methylome ^27^. This indicates that the paternal effect is of high importance for developing progeny, which should be carefully considered whenever specific sperm management protocols are designed. A recent study shows that epigenetic modifications of sperm samples subjected to cryopreservation are challenging to identify even with modern analytical tools. However, the same study reveals that highly motile sperm following cryopreservation have lower differentially methylated cytosines (DMCs) ^28^. This suggests that within the same sperm sample, there are various populations of cells with potentially different epigenetic states, which may have a direct effect on the expression of genes along embryonic development in embryos fertilized with cryopreserved sperm. Therefore, cryopreservation does not necessarily affect the methylation status (or only affects it mildly) directly in the cell but rather induces irreversible changes to the portion (subpopulation) of sperm cells with the lowest cryo-resistance, possibly also affecting the methylation profile inherited by the progeny. In this way, cryopreservation may influence the transcriptomic profile of the developing embryo by serving as a selection tool for only the most robust sperm cells. It should be pointed out that, in this way, only genes under direct paternal control will be significantly affected, which can be certainly identified in the developing progeny following a controlled fertilization operation. Thus, we were eager to compare the transcriptomic portrait between progeny obtained with the use of either fresh or cryopreserved semen. Modern transcriptomics encompasses an understanding of the RNA repertoire present in the cell along with transcription patterns, expression levels, functions, locations, trafficking, and degradation^29^. Transcriptomics is a robust, sensitive, and high-throughput technique that is typically used to identify differentially expressed genes (DEGs) between cells and tissues ^30^. The data set obtained can be further used to evaluate the function of these DEGs (among others, gene ontology). In the present study, we carried out RNA sequencing (RNA-Seq) of RNA obtained from freshly hatched larvae (at the mouth opening stage) to identify the processes being modulated/affected in the progeny by the usage of cryopreserved sperm for fertilization in Eurasian perch (*Perca fluviatilis*), which is a model for percid fishes, an important group of commercially relevant aquaculture freshwater species. The strength of the present study is its importance around the integration of information on zootechnical performance of the progeny and the transcriptomic profile/repertoire obtained from the whole organism. After all, combining transcriptomics data and associated phenotypic characteristics observed during advanced zootechnical exploration is an excellent approach to link genotype-phenotype relationships ^31^.

Eurasian perch is a commercially relevant species farmed in recirculating aquaculture systems (RASs), attaining the 4^th^ level of domestication thus far ^32^. In the last 20 years, it was found to be an excellent model for studies on embryonic development ^33^, reproduction ^34^, domestication processes ^35,36^ and circadian rhythms ^37^. In addition, the larvae of this species can be utilized as a complete organism to sequence their RNA repertoire, given their size and developmental advancement. At their mouth opening stage, they are self-sustainable organisms with the ability to adapt to different environments. More importantly, at this stage, they are not yet affected by any human intervention ^36^. In addition, a Eurasian-perch specific, highly standardized sperm cryopreservation procedure was developed by ^38^, enabling the maintenance of high fertilization success with the use of cryopreserved semen. This has brought the tool, enabling much more feasible and sophisticated selective breeding procedures in this species. However, until now, neither molecular nor zootechnical consequences have been investigated following the usage of cryopreserved sperm for the creation of a new generation in this species. In addition, prominently, any consequences passed on to the progeny from sperm subjected to these challenging procedures would be the ones directly linked with a contribution of a male to the overall phenotype of the progeny. This also includes distinguishing paternal effect genes, an important aspect in developmental biology that is highly difficult to identify. Therefore, in our study, we aimed to explore zootechnical and transcriptomic consequences in larvae resulting from the application of sperm cryopreservation technology. Controlled reproduction of Eurasian perch followed by examination of zootechnical performance became a kind of proxy for understanding physiological alterations in progeny, revealing paternally controlled genes.

## Results

Cryopreservation resulted in a significant decrease in all tested sperm motility parameters compared to fresh semen, except linearity (**Figure 1**). However, fertilization and developmental rates were not affected by cryopreservation. Additional analysis of sperm motility parameters in relation to fertilization rate did not reveal any significant correlation (see **Supplementary file 1**).

**Figure 1:**
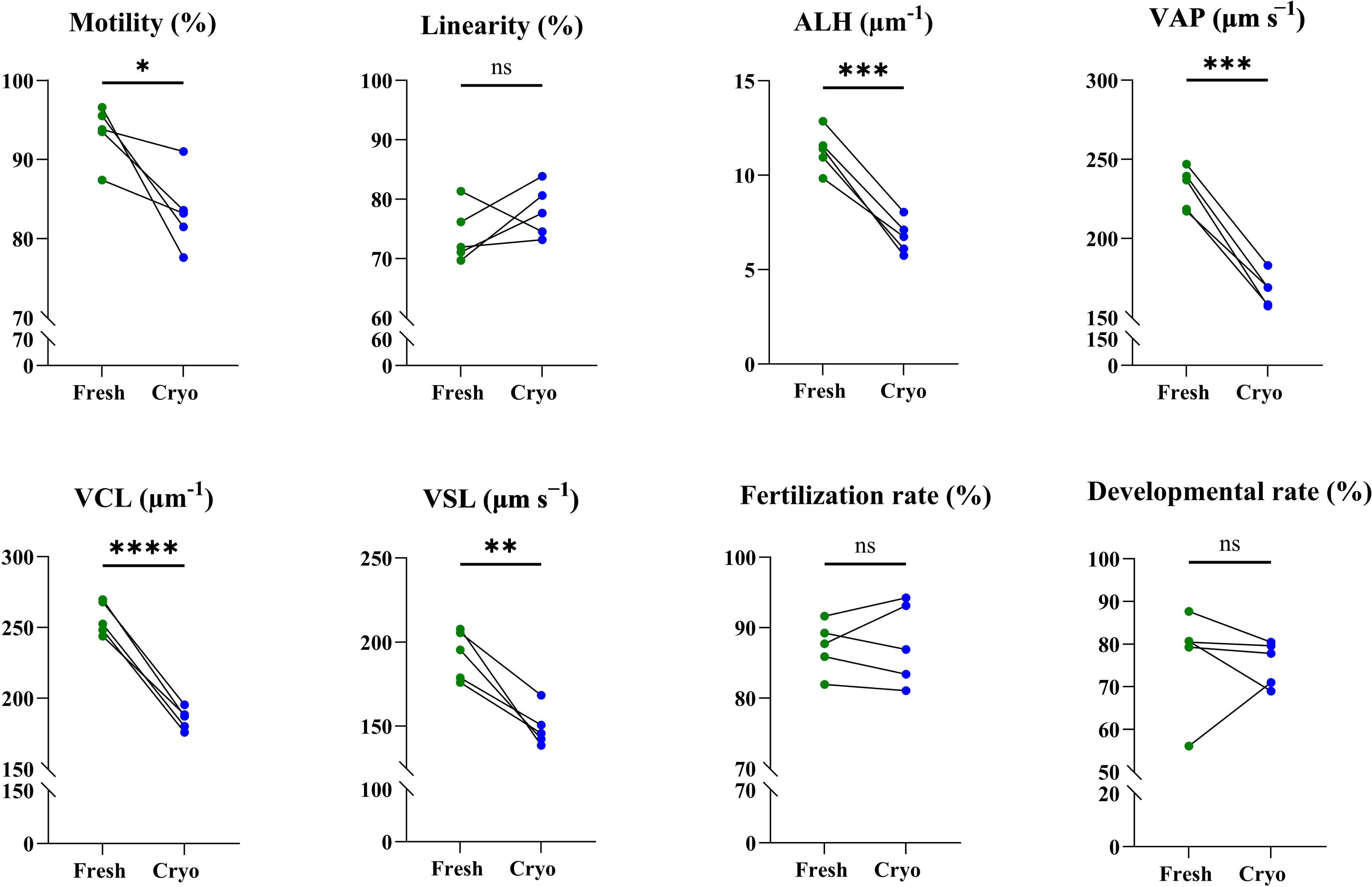
Sperm motility parameters, fertilization and developmental rate between Fresh and Cryopreserved (Cryo) Eurasian perch semen (n=5). The results for statistical analysis are presented as follows: ns – nonsignificant, * – p <0.05, ** – p <0.01, *** – p <0.001, **** – p <0.0001.) ALH-amplitude of lateral head displacement; VAP-average path velocity VCL-curvilinear velocity VSL-straight line velocity.

### Zootechnical parameters

No significant differences in deformity rate, SBIE rate, TL (both at mouth opening and weaning stages) or mortality were recorded between the Fresh and Cryo groups. However, a significantly higher WBW of the larvae from the Cryo group at the weaning stage was detected (**Figure 2**) compared to the Fresh group.

**Figure 2:**
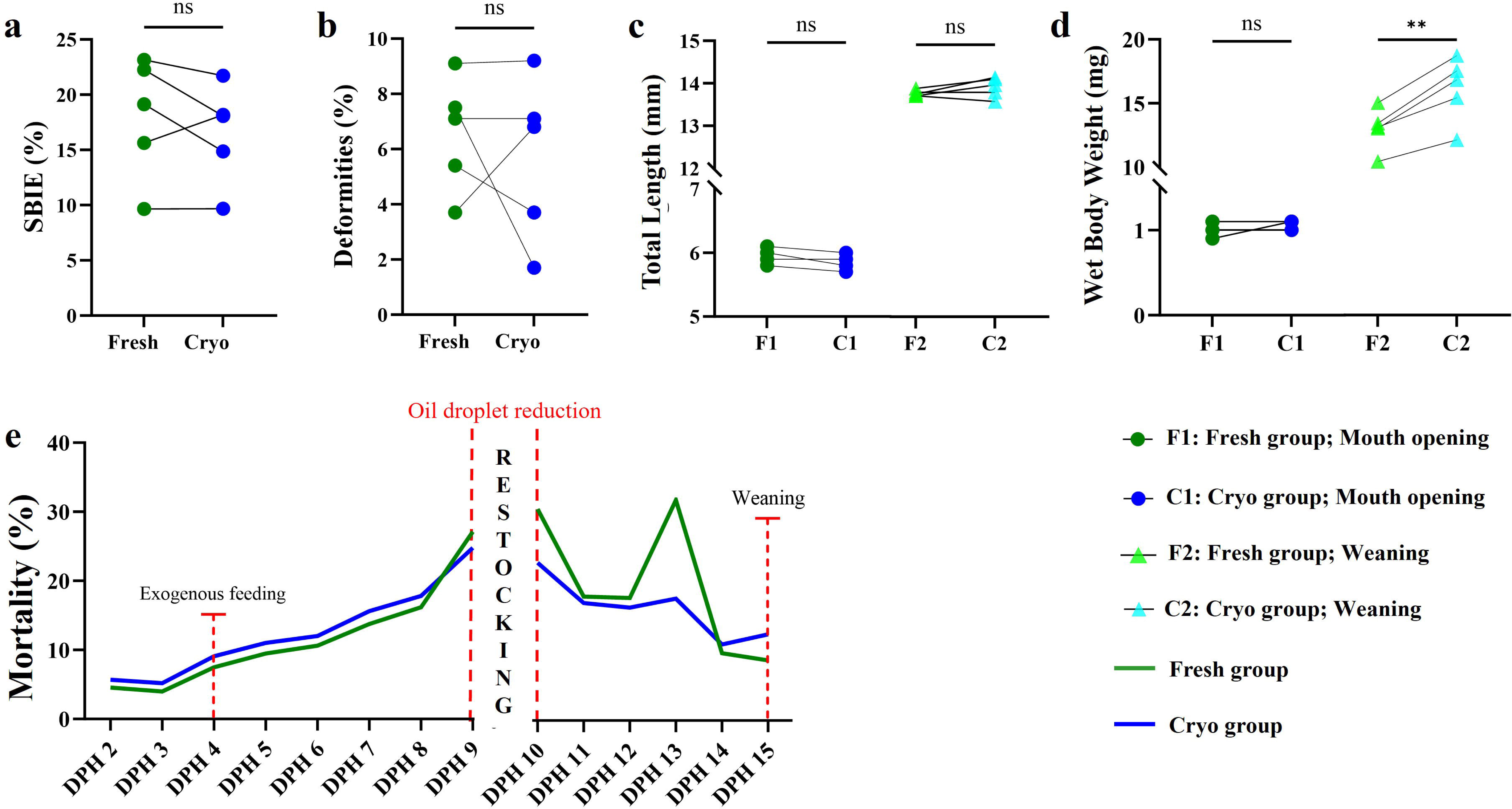
Zootechnical performance larvae obtained with the use of Fresh and Cryopreserved semen of Eurasian perch (n=5). **a**-Deformity rate at mouth opening stage (%) **b**-Swim Bladder Inflation effectiveness (SBIE, %); **c**-lengths of larvae at mouth opening and at weaning stages (mm); **d**-weights of larvae at mouth opening and weaning stages (mg); **e**-Cumulative mortality of larvae over the larviculture period, before and after restocking (only eating larvae). The results for statistical analysis are presented as follows: ns – nonsignificant, ** – p <0.01).

### Differentially Expressed Genes (DEGs)

Analysis of the transcriptomic data enabled the identification of 11 DEGs (**Figure 3**) between the Fresh and Cryo groups. The only gene with higher expression in the Fresh group was *tgfbi*, while the remaining 10 genes had higher expression in the Cryo group.

**Figure 3:**
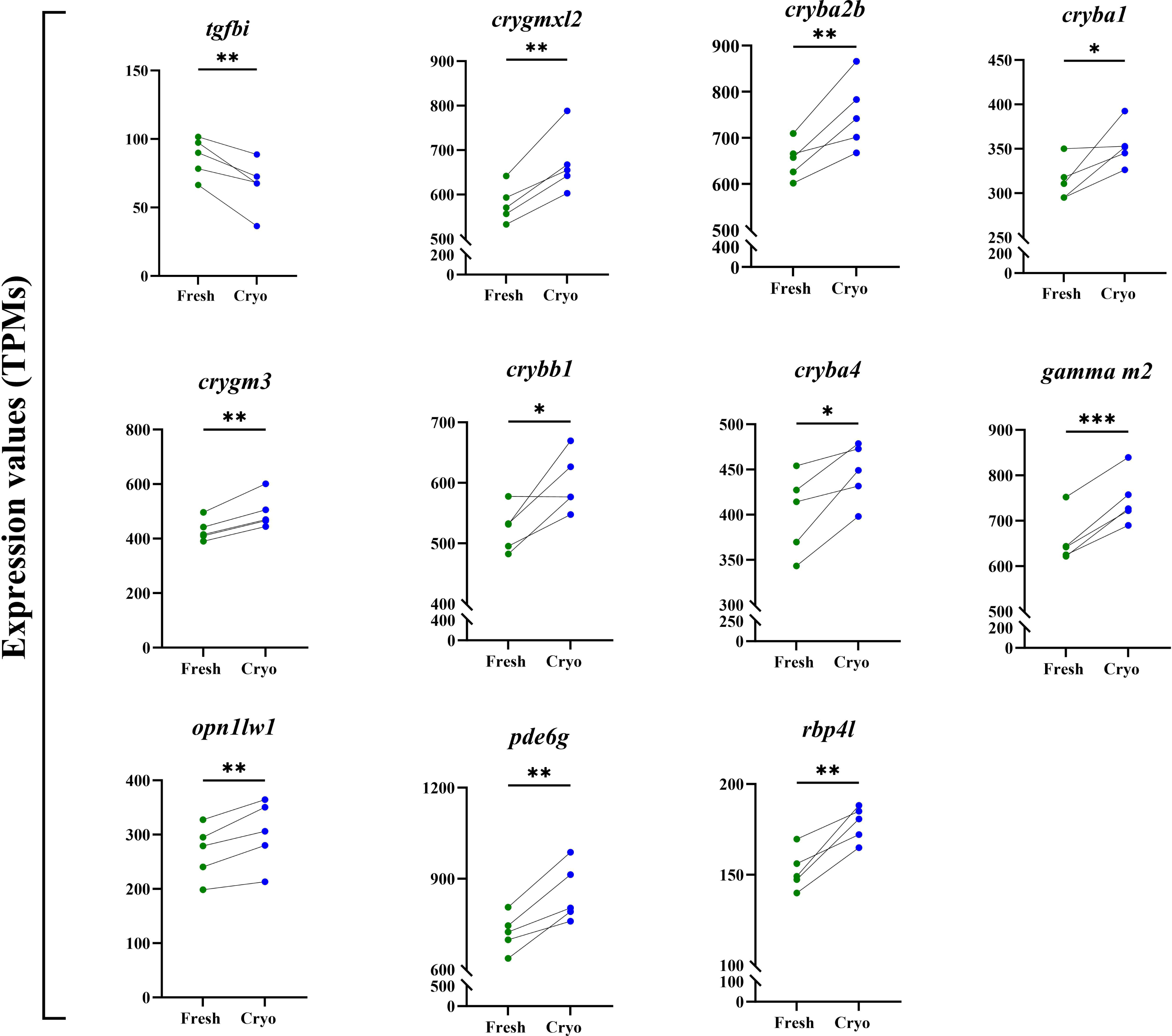
Differntially Expressed Genes (DEGs). Expression levels of DEGs identified after transcriptomic analysis of larvae at MO stage between Fresh and Cryo group (n=5). The results for statistical analysis are presented as follows:, * – p <0.05, ** – p <0.01, *** – p <0.001).

Functional analysis of identified DEGs suggested common functions in most of them. For instance, *crygmxl2*, *cryba2b*, *cryba4*, *crybb1*, *cryba1* and *crygm3* belong to the *Crystallin* family of genes, which have a major role in early embryonic eye lens development ^39^. Clustering analysis of the most enriched gene ontology terms revealed common functions related to eye development, since the remaining genes (*pde6g*, *opn1lw1*, *rbp4l* and *tgfbi*) were found to be responsible for functions of the eyes, such as photorecptors, photoperiodism, and camera-type eyes (**Figure 4**).

**Figure 4:**
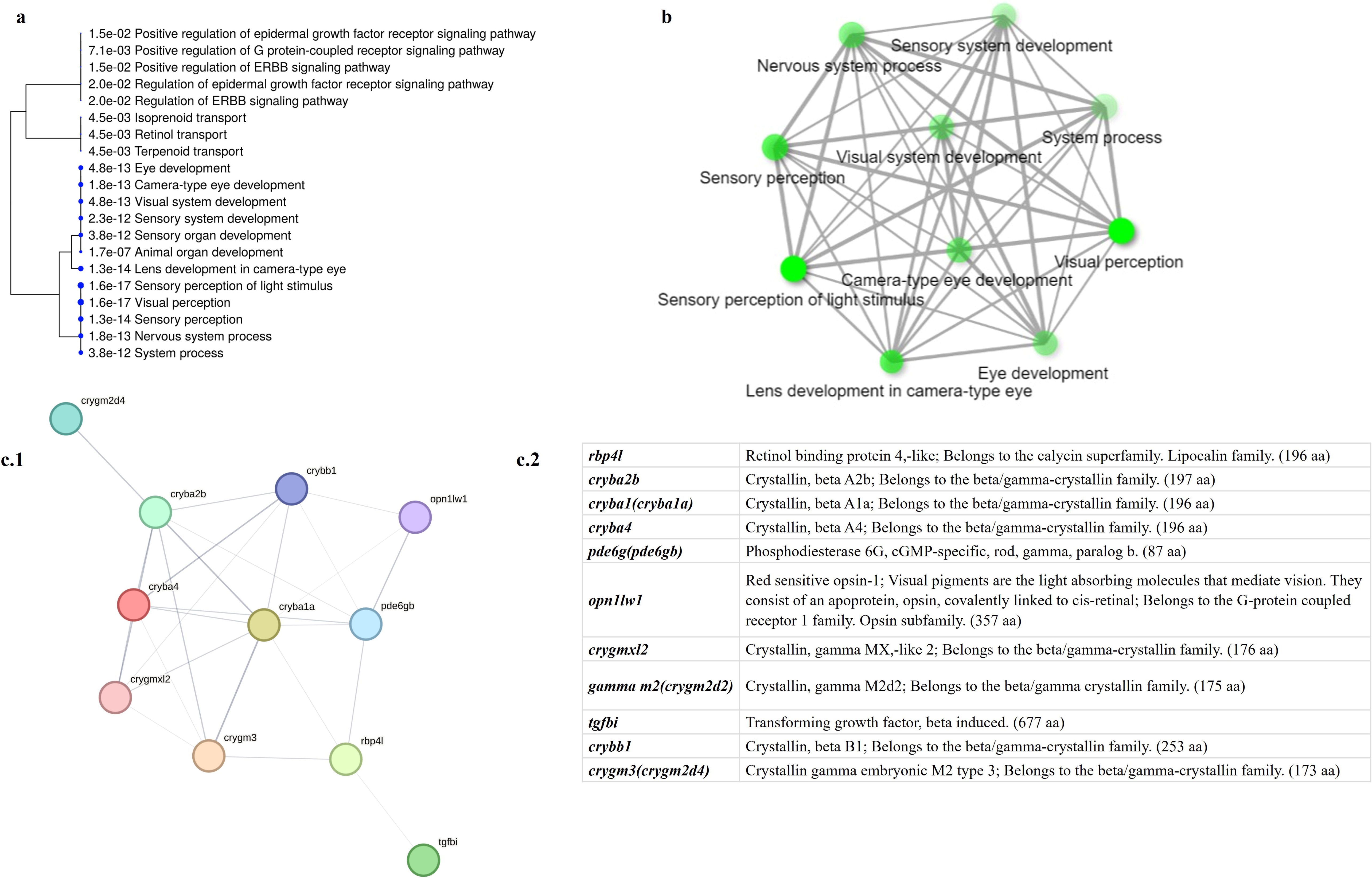
*in silico* analysis of DEGs. **a.** A hierarchical clustering tree summarizes the correlation among significant pathways based on the gene ontologies. Here 20 most enriched biological processes were clustered, bigger dots indicating more significant p-values. **b.** Relationship between 10 most enriched biological processes. Two pathways are connected if they share 20% or more genes. Darker nodes are more significantly enriched gene sets. Bigger nodes represent larger gene sets. Thicker edges represent more overlapped genes. **c.1** STRING-db chart showing proteins (encoded by DEGs) interactions and; **c.2** name of the proteins (bracketed protein names are the isoforms of the same proteins in *D. rerio*).

### qPCR validation of genes

Out of 11 DEGs, in 8 of them, the expression values at both mouth opening (MO) and weaning did not differ between the Fresh and Cryo groups based on the RTLJqPCR results (**Figure 5**). Moreover, in all these genes, the expression levels decreased with age, and there were no significant differences between the Fresh and Cryo groups. Among the validated DEGs, 3 (*pde6g*, *opn1lw1* and *rbp4l*) were confirmed to have higher expression in the Cryo group (p<0.05) (**Figure 5**) than in the Fresh group at the mouth opening stage. It should be emphasized that for these three genes, similar levels of expression between the Fresh and Cryo groups at the end of the experiment (at weaning) were observed.

**Figure 5:**
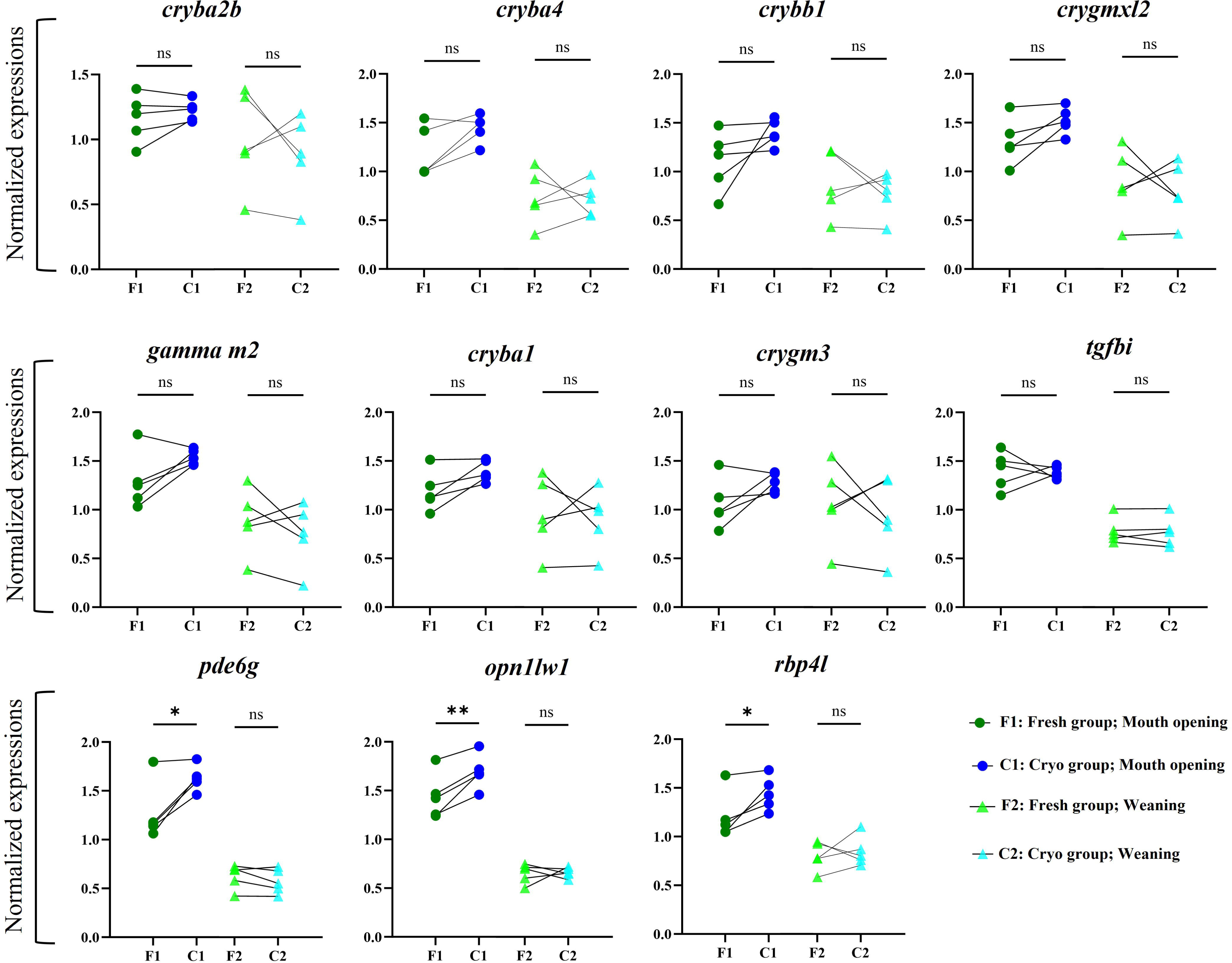
qPCRs. Normalized expression values obtained after qPCR (genes differentially expressed between Cryo group and Fresh group; n=5 for each group. The results for statistical analysis are presented as follows: ns – nonsignificant, * – p <0.05, ** – p <0.01.

### Verification of DEGs as paternal-effect genes

To add to the proof for these genes’ origin as a paternal effect, we checked for their presence in unfertilized (UF) eggs of some fish species for which data on the transcriptomic profile of UF eggs were available (**Figure 6**). While we did not observe any particular pattern among the species or genes, it is still evident that they are mostly not maternal genes (not present in the UFeggs), and we hypothesize that their expression starts after zygotic genome activation. To check whether the expression of the obtained DEGs started after zygotic genome activation, we checked their expression level during embryonic and early larval development in zebrafish (**Figure 7**). This allowed us to confirm that these genes are expressed long after ZGA, which additionally supports the hypothesis that their expression is under the control of males.

**Figure 6:**
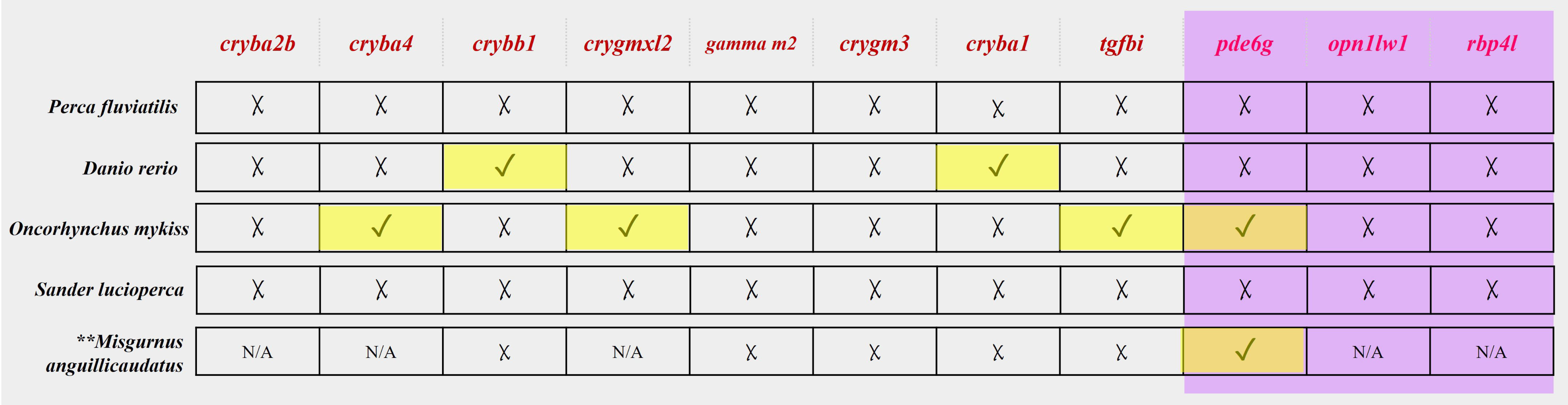
DEGs in Un-fertilized eggs. A pictograph for the presence of our candidate genes and genes validated by qPCR (written in pink and shaded in purple, respectively) in UF eggs in *P. fluviatilis*, *D. rerio*, *O. mykiss*, *S. lucioperca*, *M. anguillicaudatus*. (Reference UF eggs sequencing done as described earlier for *P. fluviatilis*, from PhyloFish and Expression atlas (TPMs >0.5 considered); ** Results based on TMM counts ^78^; N/A stands for data not available) represents presence while X represents absence of a particular gene in that species.

**Figure 7:**
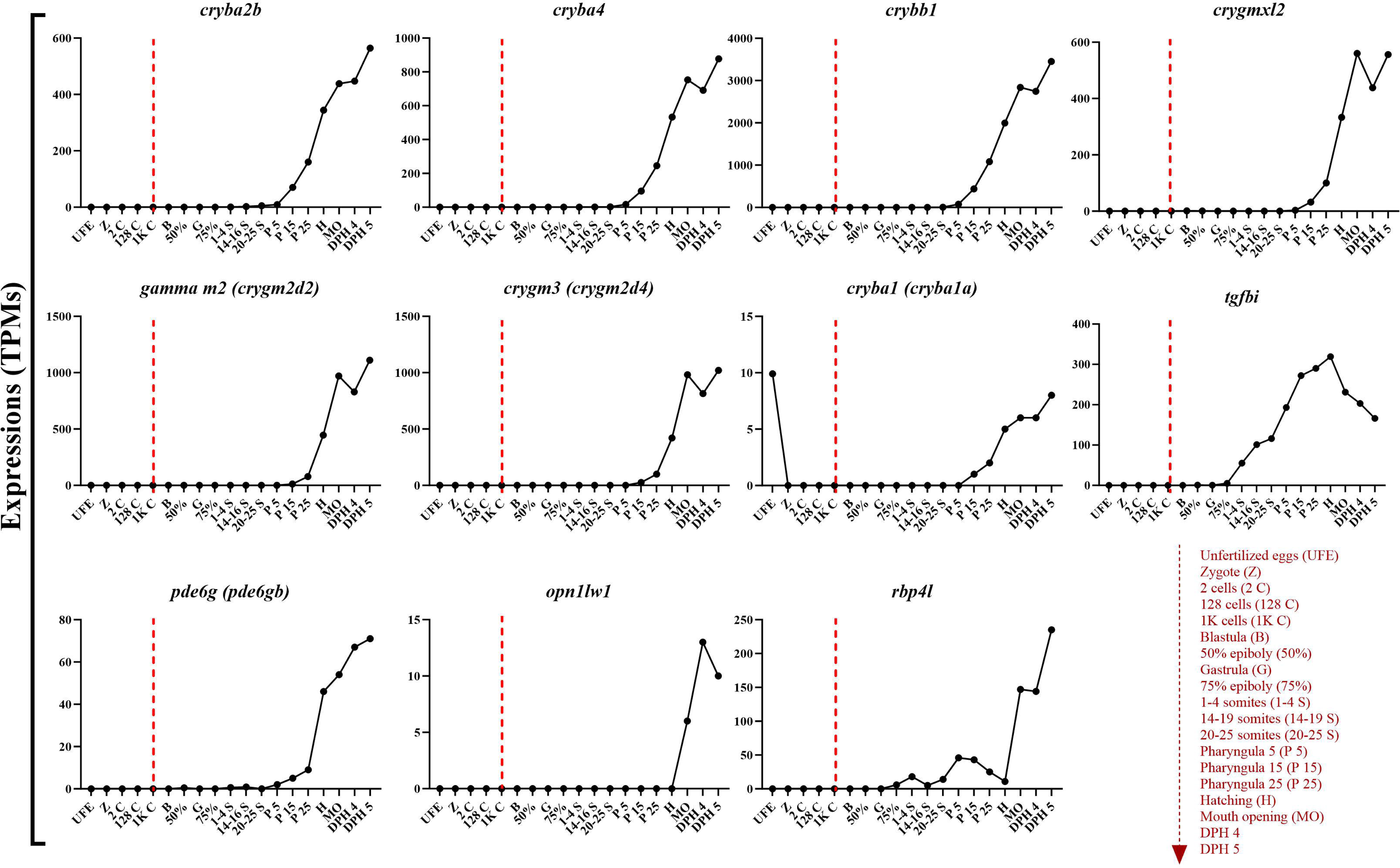
*D. rerio.* Expression level of identified paternal-effect genes along the zebrafish early development (data extracted from White et al.^77^). Abbreviations used on the x axes are explained on the right-bottom with the broken arrow indicating time course. Broken lines on each graph indicate the moment of zygotic genome activation.

### Tissue distribution of the DEGs

Given the literature background and the results of our experiment, where such important genes are differentially expressed, we wondered whether they are finite to just visual-sensory metabolic pathways or whether they have more to contribute to organism development. In this pursuit, we conducted *in silico* analysis to determine the tissue distribution of these genes across the evolutionarily distinct taxa (**Supplementary file 2**). We found that the expression of these genes was not limited to the eyes and/or brain, which could be expected for the genes directly linked with the visual system. These genes in the evolutionarily blind species cave Mexican tetra (*Astyanax mexicanus*), which has adapted to the cave environment, were also found to carry these genes in various tissues ^40,41^.

## Discussion

It is very commonly assumed in fish reproductive biology research that the most of the variation in ELH in fishes are due to maternity. However, over the years research is adding up to this fact that both the mother and the father contribute to progeny quality ^8^. The present study was aimed towards examining paternal effects on ELH traits in Eurasian perch by fertilizing eggs of individual females with either fresh or cryopreserved semen from the same male. As we assumed cryopreservation has been proven to be a challenge test for the sperm and was found to have modulatory effect on structure and functionality of diffferent spermatozoa populations. Such approach allowed us to identfy novel paternal-effect genes and led us to be the first ones to shed light on the development of visual system being under the direct control of male in fishes.

Results of our study indcates that only the cryo-resistent cells remained viable after thawing and became carriers of genetic and non-genetic information to pass on to the next generation. It has been demonstrated in rainbow trout that sperm cryopreservation did not affect fertilization rates ^28^, and no effects on development and survival during the embryo stage. However, fertilization of eggs using cryopreserved sperm led to significantly reduced larval growth after hatching ^24^. This is in contrast to our study, where after using cryopreserved semen for fertilization of eggs it resulted in increased WBW of the larvae compared to larvae obtained with fresh semen at the end of our experiment. This allows us to hypothesize, that in Eurasian perch cyopreservation-induced changes are causing permanent alterations to the cryosensitive subpopulation of sperm cells which then become non-functional and are not participating in fertilization.

In the last decades, non-genetic inheritance and transgenerational inheritance is being studied profoundly ^42,43^. Until now the paternal contribution in fishes has been shown to be associated with the methylation pattern of the genome which is transferred to the progeny ^44^. The overexpression of the identified paternal-effect genes (in Cryo group) in our study suggest that cryo-resistant sperm cells had lower methylation state what allowed us to observe higher expression of these genes in the progeny, what is in line with the suggestions of El Kamouh *et al* ^28^. This brings us closer to the hypothesis that the sperm yielding hypermethylation of these genes are the ones possibly being cryo-sensitive leading to losing their fertilizing capacit. Of course, the confirmation and possible understanding of the exact mechanisms are to be elucidated in the future, but at this point this seems to be the most rational explanation of the phenomenon observed and overexpression of the paternal-effect genes identified in our study.

We observed no significant differences in larval performances at their early life stages from the zootechnical point of view except for one important trait of the offspring, being the WBW, recorded to be higher at the end of the growth trial in Cryo group. This is contradictory to other studies, where fish obtained using cryopreserved sperm for fertilization were characterized by lower zootechnical performance ^24^. It should be underlined, that from among all the DEGs identified in our study, 10 genes were related to the visual system development. This allows us to hypothesize that visual system aids the development of visual ograns in Cryo group by upregulation of these genes and, consequently, facilitating catching their prey faster and more efficiently. As a result, fish from Cryo group were getting higher in weight by certain age. However, this hypothesis should be critically tested during a specifically designed future study.

In this study, we have demonstrated that fertilization of eggs with cryopreserved semen resulted in overexpression of genes related with the eye development. Most of these genes (*crygmxl2*, *cryba2b*, *cryba4*, *crybb1*, *cryba1* and *crygm3)* belong to the crystallin superfamily of genes with highly abundant proteins in vertebrate lineages. They are anciently identified in vertebrates and nonchordates, as α-, β-, and γ-crystallins as main sub-families ^45^. Underwater, the lens alone provides almost all the focusing power in fish, while in terrestrial species, the cornea provides most focusing power and the lens is mainly used for fine control of image formation ^46^. Certain orthologs of *cryba2b* and *crygmxl2* are directly involved in lens formation. This was demonstrated by ^47^, when they used zebrafish model and clustered regularly interspaced short palindromic repeats (CRISPR)/Cas9 technology to lose the function of lens developing gene regulatory networks (GRNs) in *foxe3* mutant. They observed smaller eyes and defective lens formation 72 hours post fertilization, after the zygotic genome activation has already begun. The Crystallin family genes which have turned out to be DEGs in our experiment does not limit only to fishes but expands to higher vertebrates like human cataract lens ^48^. It should be underlined, that our positively validated candidate genes, *pde6g*, *opn1lw1* and *rbp4l* are found as orthologs in Atlantic salmon where they were found to be responsible for ocular cataract disorders along with other genes, increasing the prevelance of vertebral deformities ^49^. The homologs of *pde6g* are been vigorously studied in lower vertebrate model species, as it is one important gene for many retinal degeneration diseases ^50^. However, the functionality of these genes has not been evaluated basing on the food intake efficiency. Our results, for the first time indicate direct linkage between DEGs responsible for eyes development with weight of larvae which is directly controlled by the male. Moreover, our *in silico* analysis of kinetics of expression along the embryonic development in zebrafish (as shown on **Figure 7**) confirmed that all the 11 genes are being extensively expressed only after 1K cell stage i.e., after the zygotic genome activation ^51^. Lack of considerable expression before that event confirms that paternally-derived molecular cargo is important in shaping the expression of these genes. Our study provides the first evidence that this important group of genes, have significant contribution to the development of eyes in fishes and other taxa, and are under paternal control.

As mentioned, major function of the DEGs identified in our study are clearly related to the development of the visual system in fishes. However, the analysis of tissue distribution (see **Supplementary file 2**) indicates that their function is somehow more complex than only to development of the eyes. This is especially evident when comparing two forms of the same species – eye-less cave Mexican tetra (*Astyanax mexicanus* - cave) and surface Mexcan tetra (*Astyanax mexicanus* – surface), where the cave form is having multi-tissue expression of these genes despite not developing eyes at all. This indicates, that paternal control over the expression of the genes identified in our study may have much more wider consequences, not limited to the formation of the eyes. This also partially explains the compensation of expression of the genes at the whole organism level observed at the end of the study. It has been reported that the eyes in Eurasian perch at hatching are constituting significant component of the entire body, as the visual system is crucial for survival of the larvae ^52^. Later the eyes are not growing anymore so rapidly as the rest of the body, especially the organs reponsible for the digestion. Having in mind, that all the DEGs were found to be expressed in at least some of the digestive-system-related organs in various species (see **Supplementary file 2**), we may suggest that the differences in expression of these genes in eyes were simply blurred by the expression of those genes in much more rapidly developing organs. This brings our attention to the fact, that our approach (i.e. studying transcriptome of the whole larvae right after hatching – at mouth opening stage) has been suitable for identification of the novel paternal-effect genes, but still more work is needed to elucidate the function of those genes having most possibly diverse roles in other organs. Besides, the expression data at the end of our experiment can not rule out the alteration of the visual system in fish coming from Fresh group, what additionally rationalizing future work focusing on the visual development as a paternally-controlled process.

### Conclusions and future aspects

In the present study, our results prove that sperm cryopreservation is a technique of undoubted importance, especially to explore paternal contribution to the progeny in Eurasian perch. Cryopreservation being used as a challenge test here, exhibited the “survival of the fittest” trait in sperm, and we could filter out paternal-effect genes. Using zootechnical and transcriptomics approach we observed that the larvae by the end of the rearing period were higher in weights most probably because of the higher expression of genes responsible for the development of eyes in the Cryo group. Here, we refer to DEGs identified, mostly responsible for the visual perception and lens formation that helped the larvae from Cryo group to feed on their prey more efficiently. We also confirmed the absence of expressions of these genes in UF eggs which means that their expression is not from a maternal genome, but is under paternal control. Furthermore, we learnt that the role of these genes is not just confined to the development of the eyes but also several other tissues of fish species varying on the phylogenetic tree, including blind Mexican tetra. With this study, we identified novel paternal-effect genes and a future direction to learn more about how does the father control gene expression patterns in the progeny. With our findings, one could deploy these genes (of course with further investigations) to be expressed in higher levels to know more from the molecular perspectives, up to assisted reproductive technologies.

### Materials and methods Ethics statement

The study was conducted according to the European and national legislation for fish welfare and approved by the Local Animal Research Ethics Committee, resolution no 5/2023. The animal study is reported in accordance with ARRIVE guidelines (https://arriveguidelines.org) for animal research.

### Broodstock management and controlled reproduction

We crossed 3 female and 6 male wild spawners (see physiological details in **Supplementary file 3**) from Mikołajki lake and Żurawia fish farm ponds, respectively. The wild fish were caught using fyke nets and transported immediately after in plastic bags filled with water and oxygen (v/v 2:1) to the research facilities of the Centre of Aquaculture and Ecological Engineering of the University of Warmia and Mazury in Olsztyn (CAEE-UWM, NE Poland). The pond-reared fish males were harvested in November, dedicated to oxygenated tanks at the Salmonid Research station of the National Inland Fisheries Research Institute in Rutki (North Poland), where they were overwintered in the flow-through system fed with riverine water (natural photothermal conditions). However, the females were caught directly during the spawning season. They were then transported in plastic bags to the CAEE-UWM for further controlled reproduction procedures. The females and males were of different origins because the capture of wild males during the spawning season is very difficult, and often these males are prone to have partially participated or even completed the spawning act before being caught. This can have a direct effect on the sperm quality obtained; thus, the males were caught earlier and overwintered. In contrast, the wild females, if overwintered in controlled conditions, tend to have lowered egg quality, affecting the quality of the larvae and therefore causing bias to the results obtained. The males and females were kept separately, according to their gamete maturity stages as recommended by ^53^ in RAS with a defined photoperiod (14 hours light:10 hours dark) and temperature (12°C±0.1) until ready for ovulation and spermiation. To promote and synchronize the spawning act of both sexes, fish were hormonally stimulated using salmon gonadoliberin analog (sGnRHa, BACHEM, Switzerland) (injection at a dose of 50 μg kg^−1^) ^53^. Sperm were collected 7 days post hormonal stimulation (which was within the optimal period of sperm collection of this species) ^54^, whereas eggs were collected between 3- and 5-days following injection depending on the maturation stage of the females ^55^. Prior to any manipulation, such as gamete collection, the fish were anesthetized in MS-222 (Argent, USA) at a dose of 150 mg L^−1^. Twelve unique families (each family reared in triplicate) were selectively created using 3 females and 6 males. More specifically, eggs from one female were divided into four portions, each fertilized with either fresh (group Fresh) or cryopreserved (group Cryo) sperm from two males, separately (as described in **Figure 8a**). Apart from gametes, other information from parents, such as total length (L_T_), caudal length (L_C_), body weight (before and after gamete stripping), body scales (for estimation of the fish’s age) and fin-clip samples, was collected (**Supplementary file 3**).

**Figure 8:**
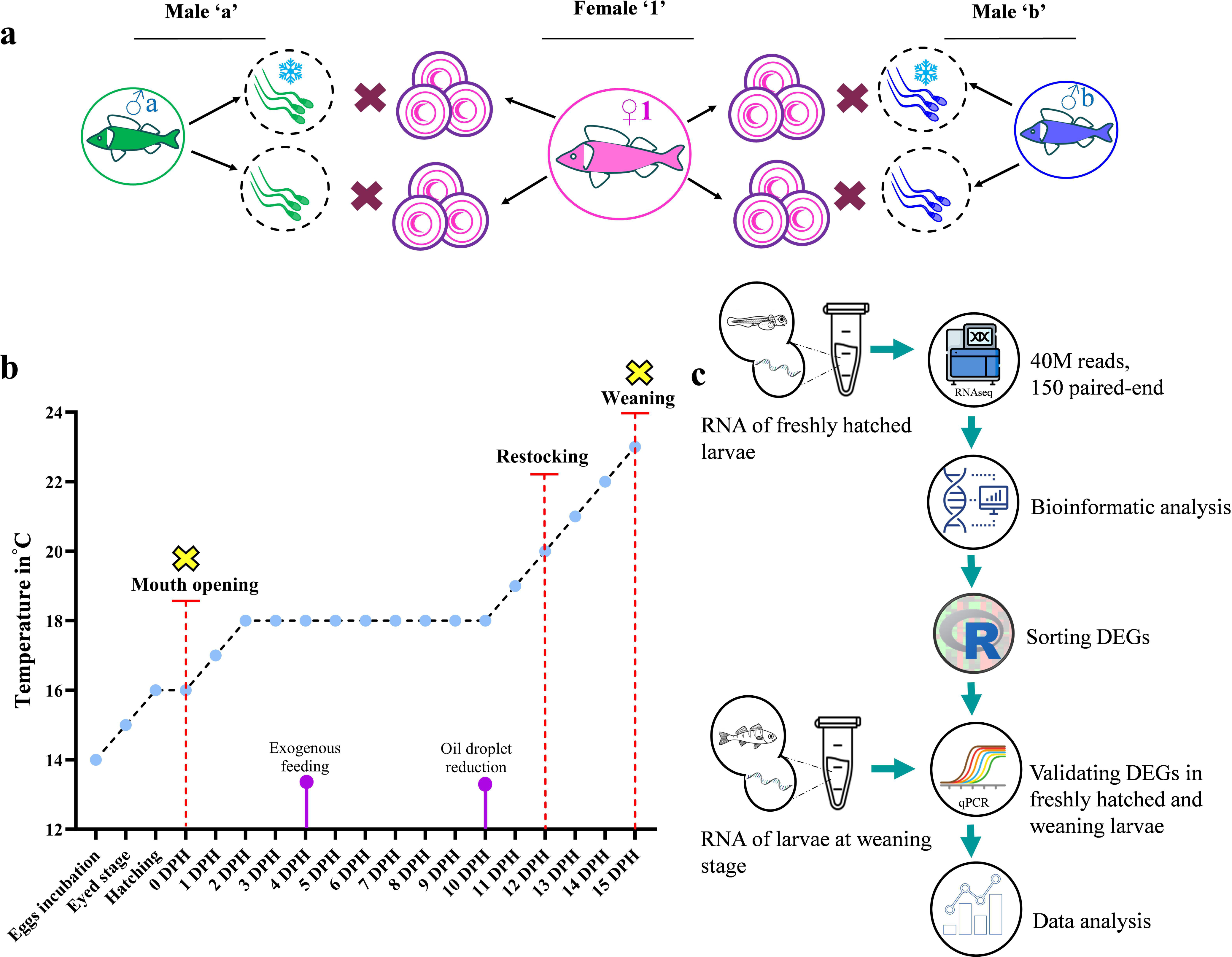
Protocol followed for the experiment. **a**: Eggs from one female was fertilized with semen from two males such that eggs from one female was divided into two portions for each male, one fertilized with cryopreserved semen (marked with a snowflake) and the other portion with fresh semen; **b**: Rearing schedule and temperature regime, sampling points (yellow crosses); **c**: RNA extraction and sequencing, followed by sorting DEGs, validations by qPCRs and data analysis.

### Sperm collection and cryopreservation

The 6 males were stripped (with gentle pressure on abdomen) for semen using a catheter (to avoid contamination of the urine or blood). After collection, each sample was kept on ice, while parameters such as sperm motility were evaluated using a computer-assisted sperm assessment (CASA) system, and the concentration was measured using NucleoCounter SP-100 (Chemometec, Allerød, Denmark) ^56^. One part of the collected semen was used for cryopreservation as described by Judycka *et al*. ^56^. The semen was diluted with extender (consisting of a concentration of 0.3 M glucose, 7.5% methanol and 25 mM KCl at 3 × 10^9^/ml spermatozoa). Semen mixed with cryoprotectants was filled into 0.5 ml plastic straws and then placed on a floating rack in liquid nitrogen vapors for 5 minutes. The other part of the semen was kept on ice to be used directly for the fertilization trials without any manipulations.

Quality parameters such as motility rate (%), linearity (LIN, %), amplitude of lateral head displacement (ALH, µm), average path velocity (VAP, µm s^-1^), curvilinear velocity (VCL µm s^-1^), and straight-line velocity (VSL µm s^-1^) were evaluated using the CASA system for both fresh and cryopreserved semen.

### Egg collection and fertilization trials

The chosen females were taken to check their oocyte maturation stages as described earlier ^55^ by catheterizing a few oocytes, exposing them in Serra’s solution (ethanol, formalin, and glacial acetic acid mixed 6:3:1 by volume) and microscopic evaluation of their maturation stages. At ovulation, eggs from females were stripped out into a clean and dry beaker as described earlier ^55^.

Each egg ribbon with an average weight of 60± 10 g was sampled into 3-5 small portions (<1 g) and weighed, and the eggs per portion were counted. In this way, we could estimate the number of eggs present per gram to aid us in dividing the ribbon into 4 equal portions (2 portions per male). Ribbon(s) from each female were further divided into equal portions (conducted the rearing in triplicate) of ∼4 g each and were used to carry out the *in vitro* fertilization ^57^. Just before fertilization, straws were thawed in a water bath at 40°C for 10 seconds and placed in an Eppendorf tube ^56^. Then, the eggs were preactivated for 30 s in hatchery water, and semen (fresh or cryopreserved) was added to the eggs at a sperm:egg ratio of 200,000:1. Then, an appropriate amount of semen was added to each egg portion, as previously calculated (separately for each egg sample at a 200,000:1 sperm:egg ratio). The eggs were then stirred for 30 seconds and washed with hatchery water after ∼10 minutes to remove excess sperm and any debris.

### Incubation of embryos

The fertilized eggs were incubated in 5 L tanks with black walls that functioned within the same RAS. Initially, before hatching, the eggs were placed on mesh (diameter of 3 mm) at a temperature of 14°C. Within 24 hours of fertilization (before embryos reached the mid-blastula transition (MBT)), ∼100 random embryos were observed under the microscope to calculate the fertilization rate. Similar counting was again performed after 3 days, while the embryonic development rate was estimated at the neurula stage (when the body of the embryo could be viewed at the animal pole). While incubating the embryos, the photoperiod was maintained at 24 L:0D, and the temperature was raised to 15°C when the embryos reached the eyed-egg stage; then, the temperature was maintained at 16°C as soon as the first hatched larvae were noticed ^36^. We started numbering the age of larvae post hatching as DPH (days post hatching), and to maintain synchronous hatching, the larvae were hatched manually (gentle actuation of the egg ribbons in a bowl). This was done by transferring the egg ribbons to bowls with water from the rearing tanks and stirring gently, and the hatched larvae were put back to their respective tanks. We carried out this operation 4-5 times. That day was named 0 DPH, and the day count had begun.

### Larviculture and advanced zootechnics

The hatched larvae in both the Fresh and Cryo groups were reared following the exact same conditions in the RAS system, along the standardized temperature and feeding regime described (**Figure 1b**) ^36^. Beginning from 0 DPH, the temperature was 16°C. At 1 DPH, the water temperature was raised by 1°C, and at 2 DPH, it was at 18°C; this temperature was kept stable up to 10 DPH. Starting from 4 DPH feeds of *Artemia* sp. nauplii *ad libitum* three times per day (first four days of feeding – micro *Artemia* cysts [SF origin], then standard size *Artemia* cysts at 260,000 nauplii per gram [GSL origin]) was insured. At 12 DPH, eating larvae were restocked equal numbers of larvae in all tanks by counting volumetrically. Here, eating larvae ensured healthy larvae to a few extents. Subsequently, 11 DPH onwards, the temperature was increased by 1°C per day until 23°C, which is considered the optimal temperature for the growth of perch larvae ^58^. After the first feeding and before the last feeding, the tanks were cleaned, and dead larvae were counted. In addition, other parameters, such as the oxygen level to 80% and ammonia concentration to <0.02 mg L^-1,^ were maintained. The experiment was conducted for the larvae only until their *Artemia* feeding phase, i.e., the experiment was terminated at the weaning stage, upon sampling.

At the mouth-opening stage (between 0 and 1 DPH, where at least 50% of larvae were found to have reached this stage) and at the time when the protocol envisaged the end of feeding, the larvae with live *Artemia nauplii* (hereinafter referred to as weaning; 15 DPH) were sampled for total length (TL, ±0.01 mm) and wet body weight (WBW, ±0.1 mg). Additionally, samples of larvae were used for extraction of RNA (whole larvae were preserved in RNAlater, SigmaLJAldrich, Germany). During the first sampling, we made sure we collected larvae at the mouth opening (MO) stage, where in-egg embryonic development has been accomplished and larvae were ready to survive in the outer environment but with minimal human intervention applied.

The total length of larvae was determined using a stereoscopic microscope (Leica, Germany). Next, wet body weight measurements using the ‘noninvasive method’^59^ were addressed using a precision laboratory scale (Ohaus, USA). For this purpose, anesthetized larvae were placed on a platform made of nylon mesh (with a mesh size of approx. 200 μm), and excess water was drained out by filter paper. This method minimized possible physical damage to very delicate larvae. Two days after oil droplet reduction, the swim bladder inflation efficiency (SBIE%) was calculated using a stereoscopic microscope by triple counting perch larvae (with and without a filled swim bladder) randomly caught from each tank on a Petri dish (in total, we determined SBIE from more than 100 larvae from each tank). Before any manipulations, we anesthetized the larvae in a solution of MS-222 (at a dose of 150 mg L^−1^).

### RNA extraction

The total RNA was extracted from snap frozen unfertilized (UF) eggs (∼50 eggs) and a pool of larvae (n=10 for larvae at mouth opening and n=4 for larvae at weaning) using a TotalRNA mini-kit (A&A Biotechnology, Poland) from unfertilized eggs of each female and larvae from each family (for both sampling stages), separately. The quantity and purity of extracted RNA were evaluated using a NanoDrop 8000 spectrophotometer (Thermo Fisher Scientific, USA). Samples showed absorbance ratios A260/280 ≥ 2.0 and A260/230 ≥ 2.2. The quality of the extracted total RNA was also evaluated using an Agilent Bioanalyzer 2100 (Agilent Technologies, USA), and all the samples presented RIN ≥ 9.0. Samples were then outsourced for RNA sequencing.

### RNA sequencing and bioinformatics

Libraries (using TruSeq stranded mRNA kit) were sequenced using Illumina’s NovaSeq 6000 with standard protocols. Overall, from each sample, more than 40 M reads were obtained, with a 150 bp paired-end sequencing mode.

### Differential analysis

The raw reads were quality controlled using FastQC software version 0.11.9 ^60^. Adapters and low-quality fragments of raw reads (average *Q*Phred score < 20) were trimmed out, and reads were clipped to equal lengths of 100 nt using the Trimmomatic tool ver. 0.40 ^61^. The resulting read sets of the analyzed samples were mapped to a reference genome *P. fluviatilis* version 11.1.104 obtained from the NCBI database ^62^ using STAR software ver. 2.7.10a ^63^ with ENCODE default options.

Transcript count data for the larval samples were filtered to have at least 5 libraries in which there were at least 5 reads. Libraries from before and after the cryopreservation process were compared using the following design: *∼ males + condition;* males standing for the 6 males followed during the experiment and condition representing before (fresh) and after cryopreservation. These analyses were performed in RStudio (version 4.1.3) using the package DESeq2 ^64^ and *ashr* for log fold-change shrinkage ^65^. Differences were considered significant when corrected p values were inferior to LJ (LJ=0.05), and we obtained 11 DEGs.

It should be emphasized that among the 6 families created and used for the entire study, for further analysis, 1 family (from the Cryo group and its counterpart in the Fresh group) was removed because the transcriptomic profile clearly differed from the remaining families and was considered an outlier (see **Supplementary file 4**).

### Gene Ontology Enrichment analysis (GOEA)

GOEA was performed using ShinyGO, version 0.77 platform ^66^ to test the overrepresentation of GO terms in a list of genes and to understand their biological significance as an effective approach ^67,68^. The 11 DEGs (namely, crystallin beta A2b (*cryba2b*); crystallin beta A4 (*cryba4*); crystallin beta B1 (*crybb1*); crystallin, gamma MX, like 2 (*crygmxl2*); phosphodiesterase 6G, cGMP specific, rod, gamma (*pde6g*); opsin 1, longwave-sensitive, 1 (*opn1lw1*); gamma-crystallin M2-like (*gamma m2*); beta-crystallin A1-like (*cryba1*); gamma-crystallin M3-like (*crygm3*); retinol binding protein 4, like (*rbp4l*); and transforming growth factor beta induced (*tgfbi*)) were fed to the ShinyGO platform, zebrafish was chosen as the best matching species; with the false detection rate (FDR) cutoff of 0.05, and 20 pathways’ network was created. A STRING-db, version 12.0 ^69^ with functional enrichment of GO biological processes was also performed to retrieve a proteinLJprotein network that also describes the distance between the linked genes.

### RT**□**qPCR validation of differentially expressed genes (DEGs)

#### Primer design

Primer pairs for all 11 DEGs along with 5 normalizing genes for RT qPCR were designed using NCBI-Primer BLAST, version 1.0.1 ^70^. The sequence that matched the best to *P. fluviatilis* was fed to Primer3Plus software version 3.3.0 ^70,71^. The best matching pairs with least possibilities to form secondary structures were chosen and checked for GC content and melting temperature (Tm) on µMelt Quartz, version 3.6.2 ^72^. The sequences of the designed primers are presented in **Supplementary file 5**.

#### qPCRs

RTLJqPCRs were performed for each gene using a Viia7 (Applied Biosystems) thermocycler. For each qPCR, 10 ng cDNA template was used along with 10 µl (A&A Biotechnology) SYBR RT PCR Master Mix (Cat. No. 2008-100), 0.5 µM forward (1 µl) and reverse (1 µl) primers, 2 µL of starter mix and PCR grade water were added to maintain a final volume of 20 µL. The reactions were performed with the following cycling conditions applied: enzyme activation for 10 minutes at 95°C followed by 40 cycles of denaturation at 95°C for 15 seconds and annealing and elongation at 60°C for 1 minute. In the analysis of each gene, a standard curve was calculated using a series of 6 twofold dilutions to determine reaction efficiency (reaction efficiencies between 85% and 110% were considered acceptable). Relative expression for each gene was normalized as the geometric mean of expression values recorded for 5 reference genes (namely, cytochrome c-like, transcript variant X1, *cycs*; tetraspanin 7, *tspan7*; ER membrane protein complex subunit 10, transcript variant X2, *emc10*; pre-mRNA-splicing factor, *syf2* and ER membrane protein complex subunit 3-like, *emc3*), which were chosen from our transcriptomic data on the basis of their stable expression levels and close-to-mean expression values in the RNA-sequencing analysis ^73^. Each reaction for real-time qPCR validation was performed in triplicate. The data were compared between the Fresh group and Cryo group (at mouth opening and weaning stages).

### In silico analysis

Several *in silico* analyses were carried out using tools such as NCBI-BLAST ^74^, Expression Atlas version 2.0 ^75^ and PhyloFish ^76^. These tools helped us to study the expression levels of our DEGs throughout the early life history in *Danio rerio* ^77^ as a reference model organism for Eurasian perch. We also performed qPCR with the RNA of snap frozen eggs of Eurasian perch used for this experiment to check for the presence of our DEGs. Along with *D. rerio*, Zebrafish, we selected a few evolutionarily close/distant species (namely, *O. mykiss*, Rainbow trout; *S. lucioperca*, Pike perch; *M. anguillicaudatus*, Dojo loach; *A. mexicanus*, surface Mexican Tetra; *A. mexicanus* cave Mexican tetra and *A. Anguilla*, European eel) from Eurasian perch to check the presence of these 11 genes. This special *in silico* analysis gave us information about the presence of these genes in different tissues in different species. This was done to get hints if a particular gene has tissue-specificity.

### Statistical analysis

The raw data from all the parameters like sperm quality parameters (in %, µm, µm s^-1^), fertilization and embryonic developmental rates (%), deformities (%), SBIE (%), TL (mm), and WBW (mg) were first fed into GraphPad (version 9.5.1) and paired t-tests (p<0.05) for each single parameter to compare between Fresh group and Cryo group were conducted. While calculating then plotting cumulative mortality (%); expression values of our DEGs (in TPMs) after sequencing and normalized expression values after real time qPCRs (mean quantity); transformation of gene replicates in TPMs for presence of genes in tissues were calculated on Microsoft Excel. However, the values were then computed on GraphPad to plot graphs after paired T-tests. All the data were tested with a significance level of 5% (significant differences were considered at p value < 0.05).

## Acknowledgements

This research was funded by the project “Transcriptomic and zootechnical exploration of parental contribution to progeny quality in Eurasian perch, *Perca fluviatilis*” granted by National Science Centre of Poland (SONATA BIS project, number UMO-2020/38/E/NZ9/00394). The authors would like to acknowledge Dr. Jarosław Król for his valuable help with logistics. Also Prof. Stefan Dobosz and Rafał Rożyński for providing facilities to maintain the fish overwinter. Additionally, the authors would like to thanks Dr. Joanna Nynca for her critical review of the draft of the manuscript.

## Authors’ contribution

**A.P.:** Conceptualization, Methodology, Validation, Formal analysis, Investigation, Resources, Data curation, Writing – Original draft, Writing – Review and Editing, Visualization. **S.J.:** Conceptualization, Methodology, Validation, Formal analysis, Investigation, Data curation, Writing – Review and Editing, Supervision, Project administration. **K.PZ.**: Conceptualization, Methodology, Investigation, Resources, Writing – Review and Editing, Supervision. **R.D.:** Investigation, Validation, Writing – Review and Editing. **S.J.:** Investigation, Validation. **J.J.:** Formal analysis, Data Curation. **T.R.A.:** Formal analysis, Data Curation. **M.B.:** Resources. **P.H.:** Resources, Data curation. **S.K.:** Resources. **D.**Ż.: Conceptualization, Methodology, Validation, Investigation, Resources, Data curation, Writing – Review and Editing, Supervision, Project administration, Funding acquisition.

## Additional information

Raw data of RNAseq used for analysis in the present study can be accessed via the NCBI SRA database (SUB13931706; restricted availability prior publication).

