## Supplementary figures and images for "Paternal-effect genes revealed through semen cryopreservation in *Perca fluviatilis*"

### Supplementary file 1

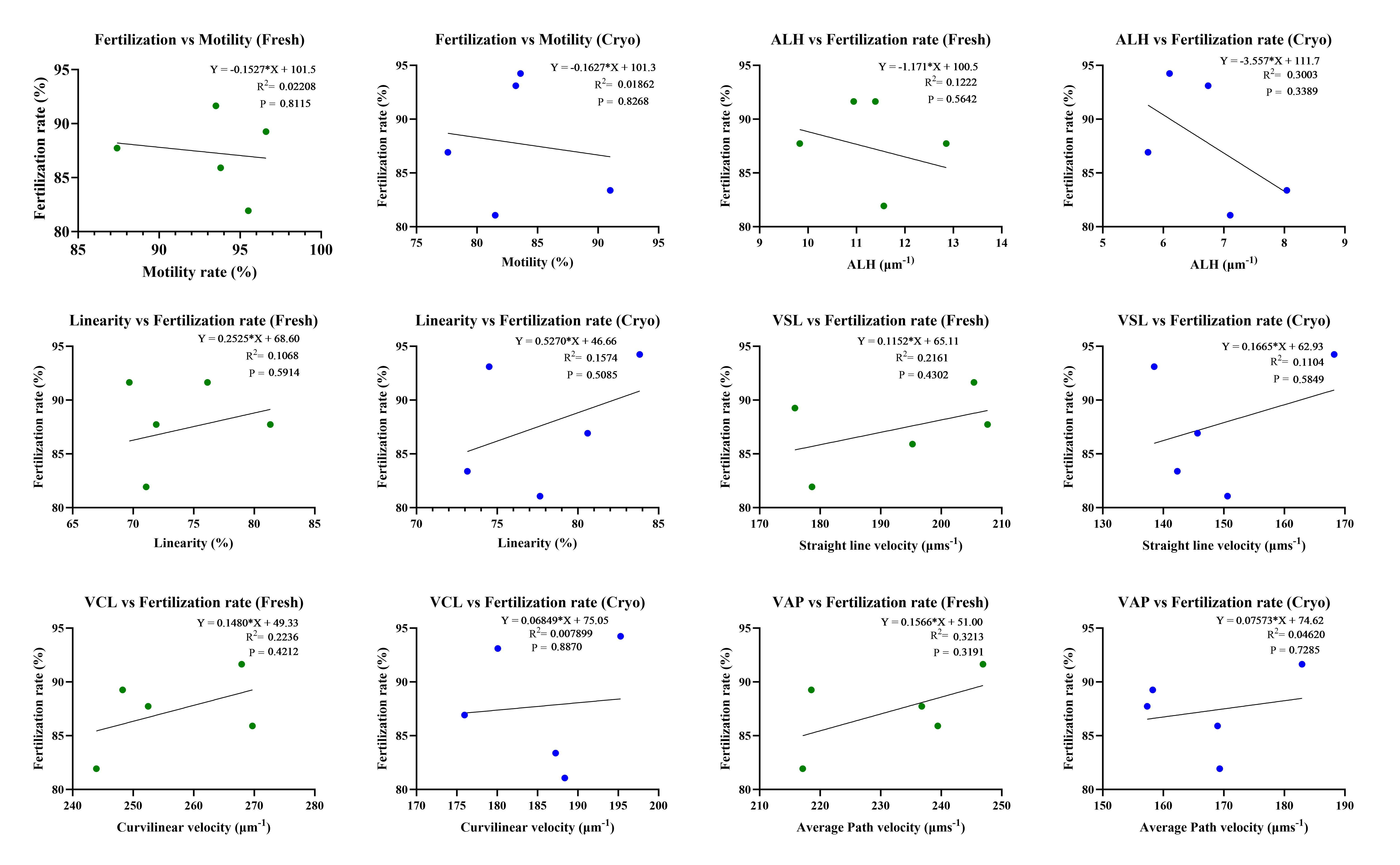

### Supplementary file 4

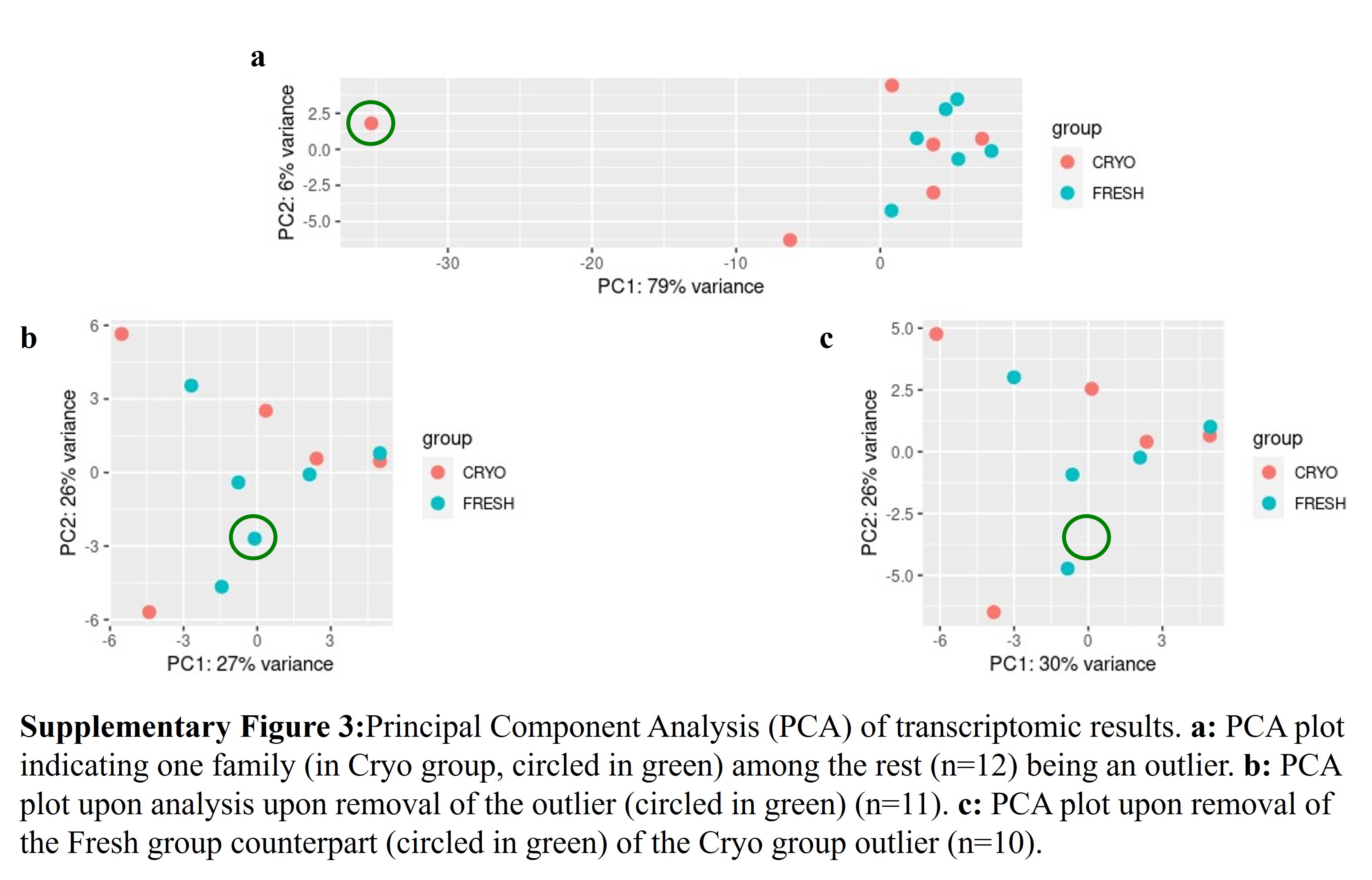
