## Supplementary file 2 for "Paternal-effect genes revealed through semen cryopreservation in *Perca fluviatilis*"

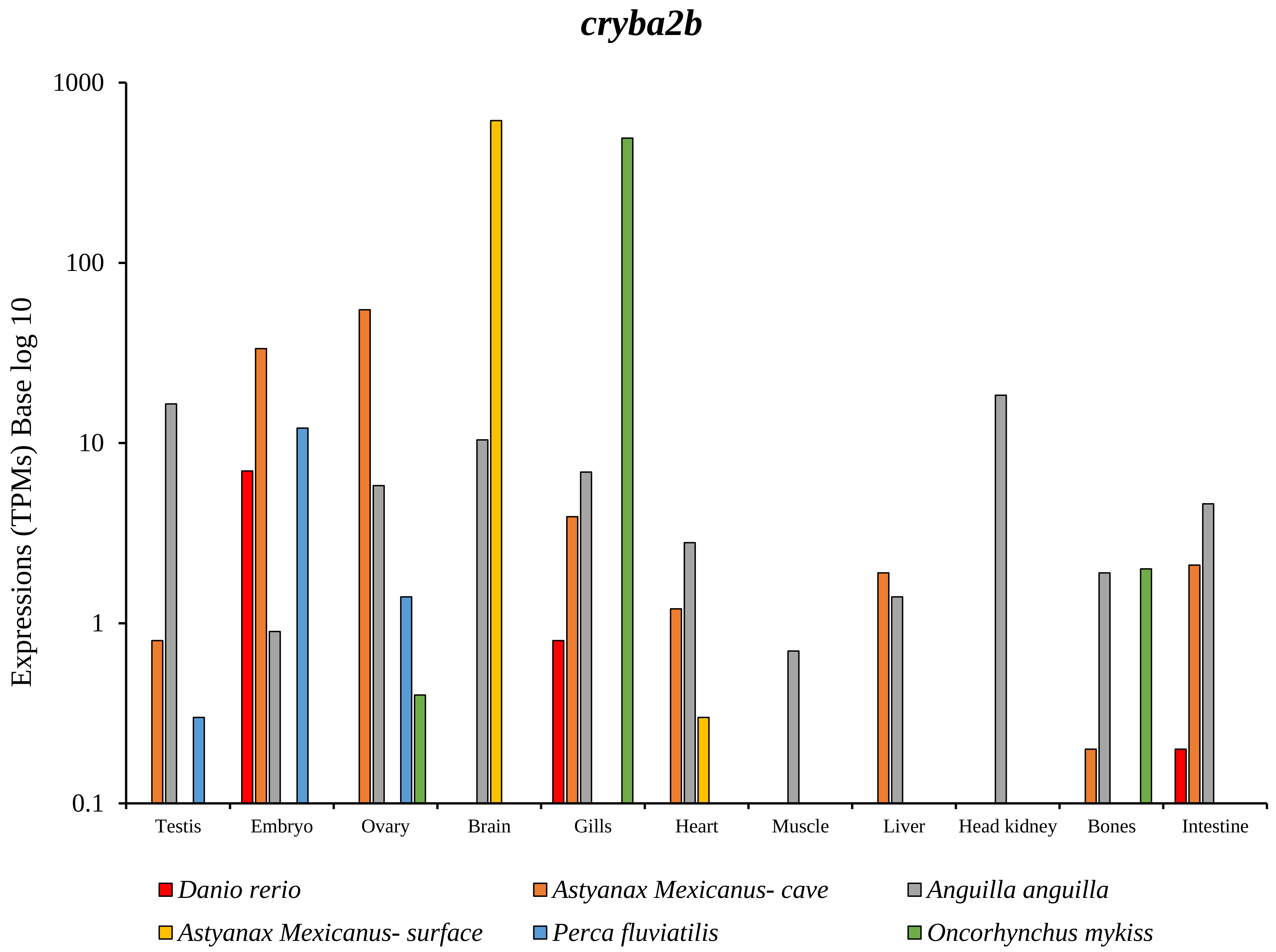
cryba2b
Expressions (TPMs) Base log 10
1000
100
10
1
0.1
Testis
Embryo
Ovary
Brain
Gills
Heart
Muscle
Liver
Head kidney
Bones
Intestine
Danio rerio
Astyanax Mexicanus- cave
Anguilla anguilla
Astyanax Mexicanus- surface
Perca fluviatilis
Oncorhynchus mykiss

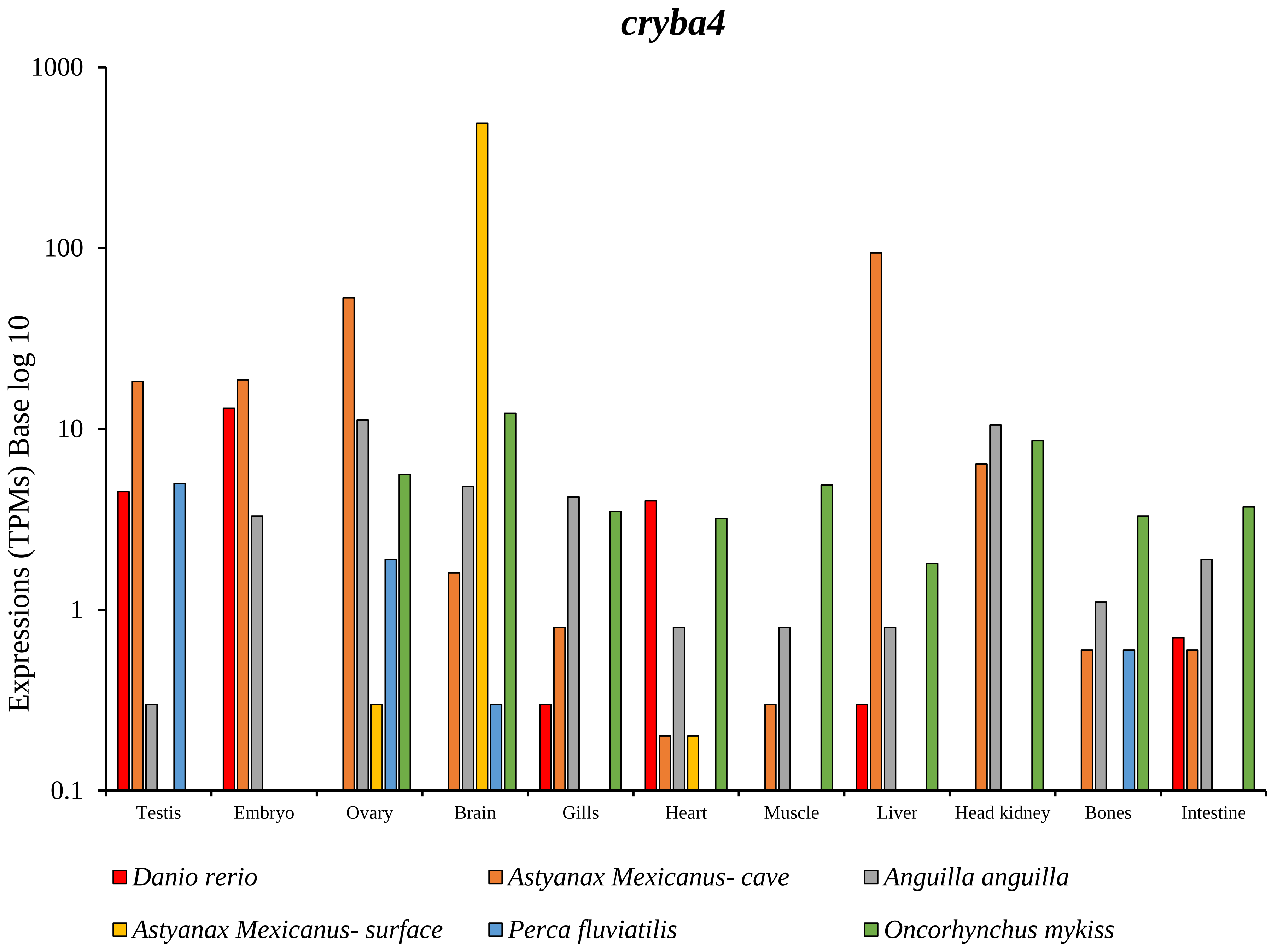
cryba4
Expressions (TPMs) Base log 10
1000
100
10
1
0.1
Testis
Embryo
Ovary
Brain
Gills
Heart
Muscle
Liver
Head kidney
Bones
Intestine
Danio rerio
Astyanax Mexicanus- cave
Anguilla anguilla
Astyanax Mexicanus- surface
Perca fluviatilis
Oncorhynchus mykiss

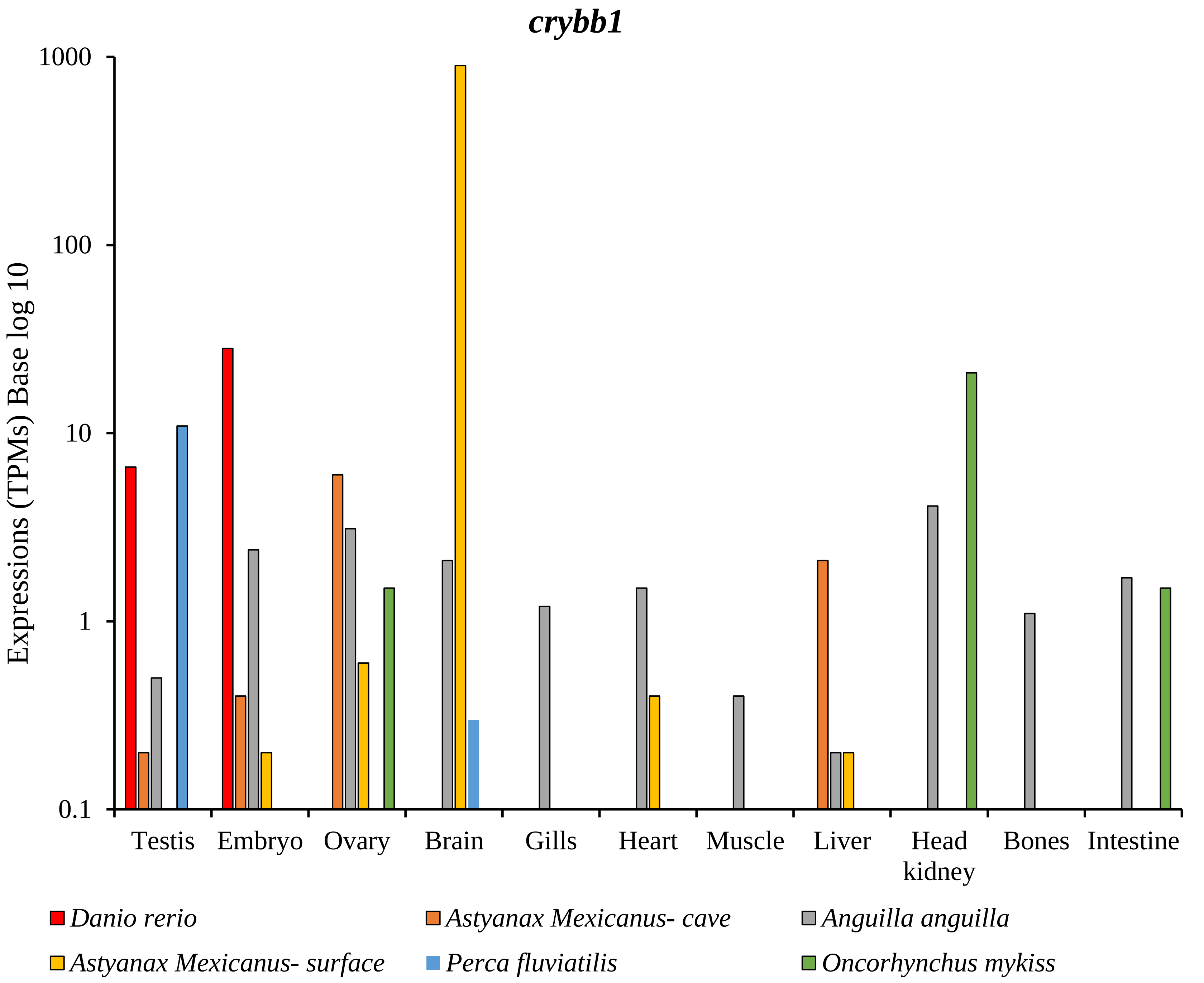

crybb1
Expressions (TPMs) Base log 10
1000
100
10
1
0.1
Testis
Embryo
Ovary
Brain
Gills
Heart
Muscle
Liver
Head kidney
Bones
Intestine
Danio rerio
Astyanax Mexicanus- cave
Anguilla anguilla
Astyanax Mexicanus- surface
Perca fluviatilis
Oncorhynchus mykiss

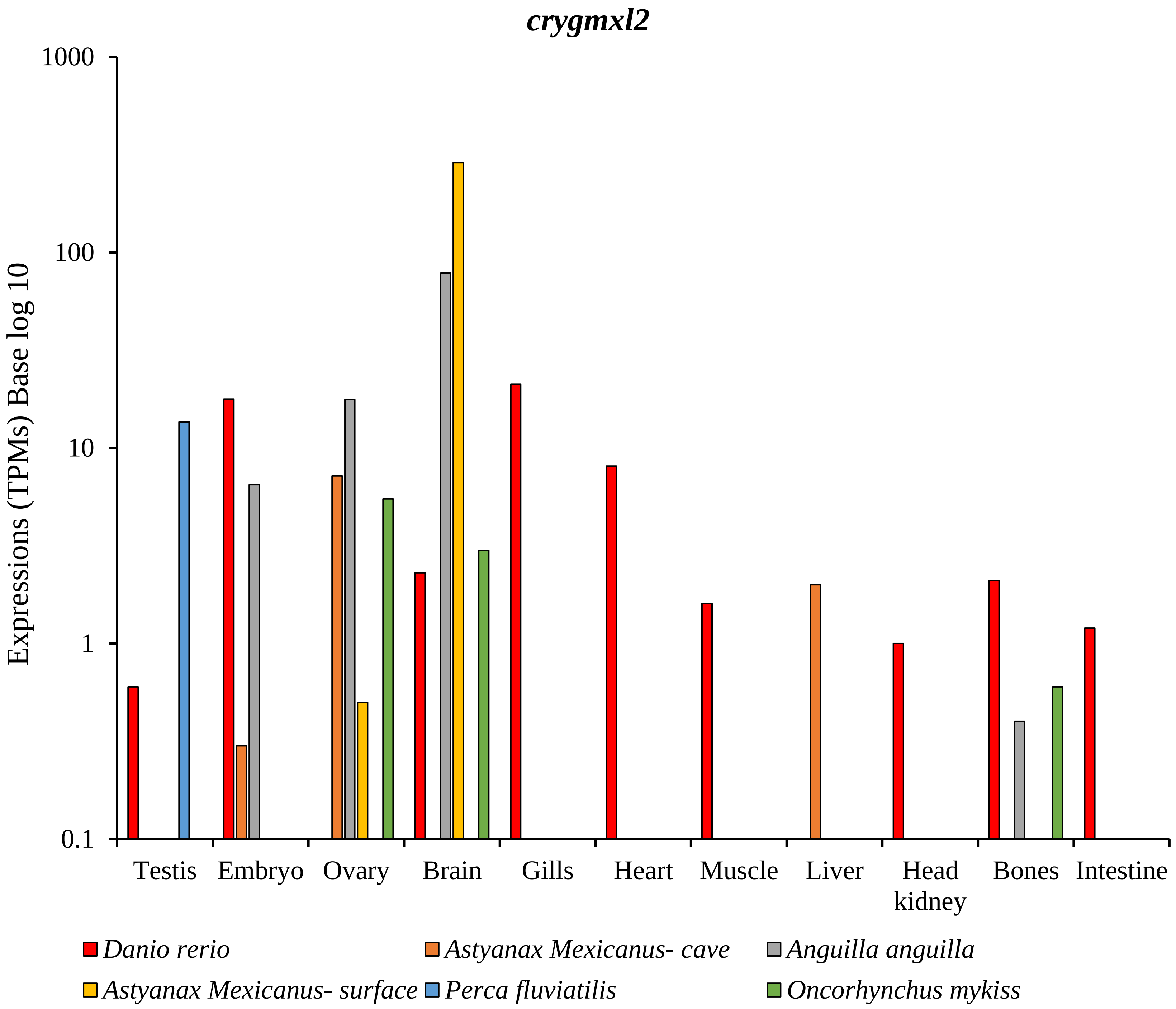
crygmxl2
Expressions (TPMs) Base log 10
1000
100
10
1
0.1
Testis
Embryo
Ovary
Brain
Gills
Heart
Muscle
Liver
Head kidney
Bones
Intestine
Danio rerio
Astyanax Mexicanus- cave
Anguilla anguilla
Astyanax Mexicanus- surface
Perca fluviatilis
Oncorhynchus mykiss

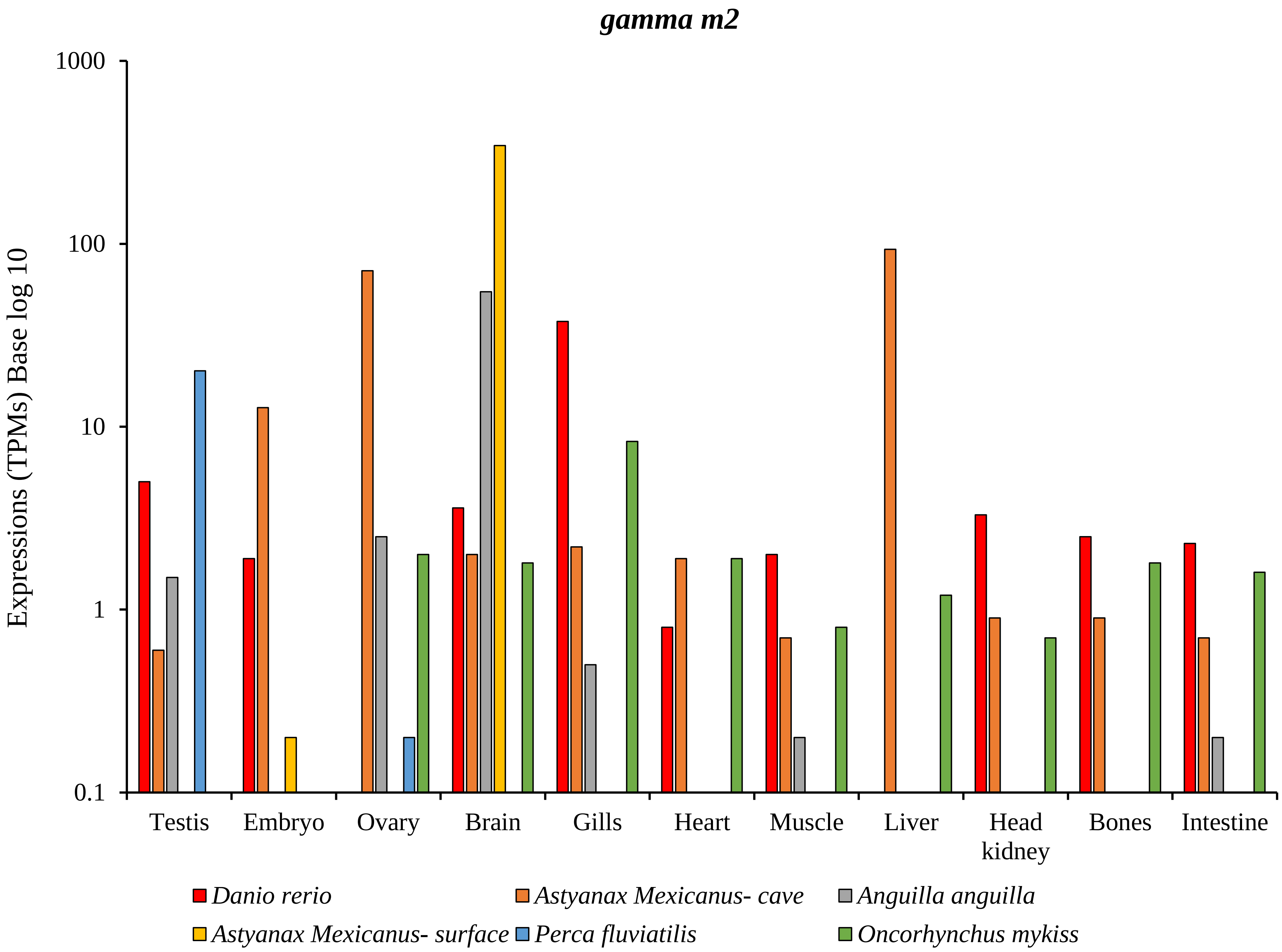

gamma m2
Expressions (TPMs) Base log 10
1000
100
10
1
0.1
Testis
Embryo
Ovary
Brain
Gills
Heart
Muscle
Liver
Head kidney
Bones
Intestine
Danio rerio
Astyanax Mexicanus- cave
Anguilla anguilla
Astyanax Mexicanus- surface
Perca fluviatilis
Oncorhynchus mykiss

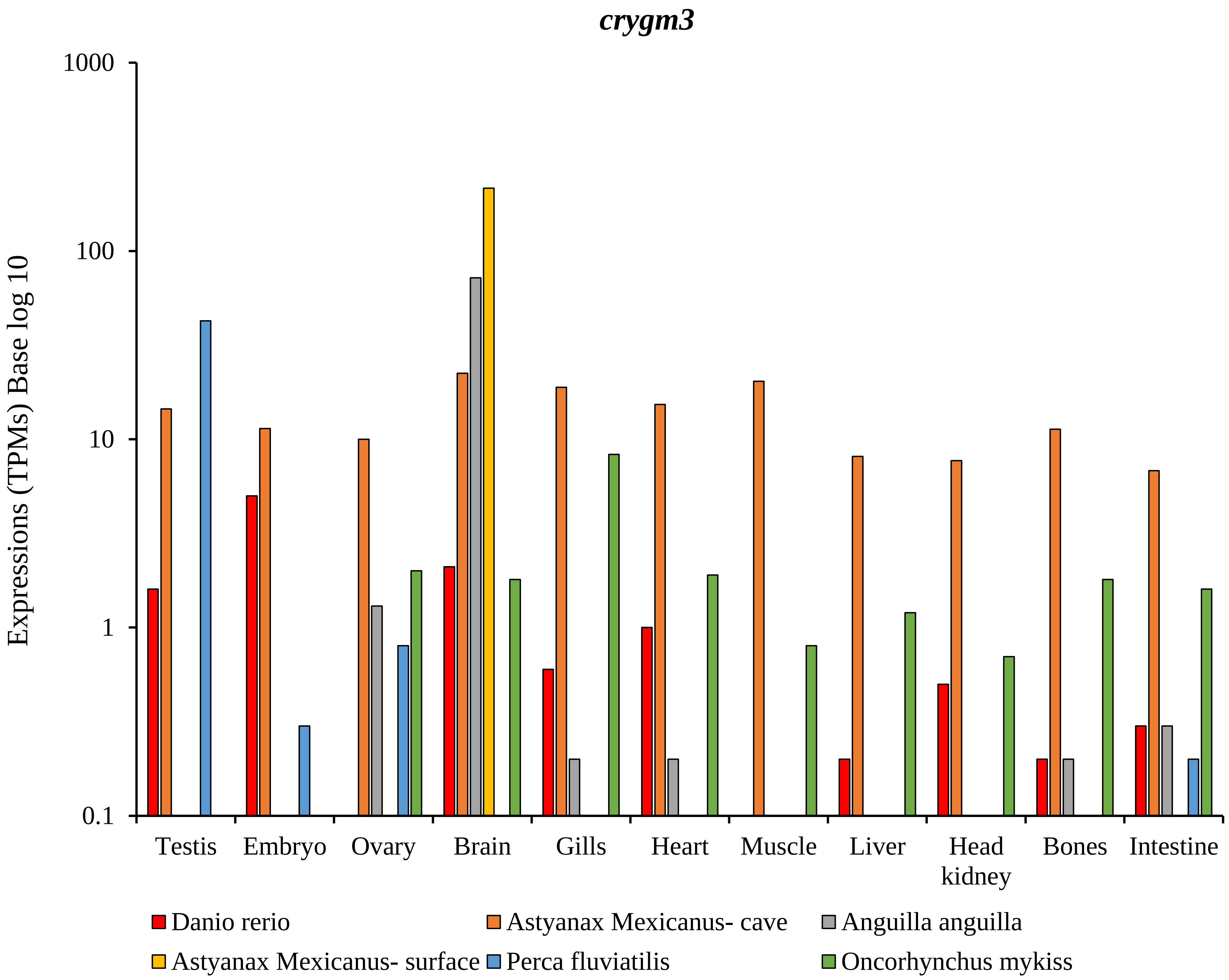

crygm3
1000
100
10
1
0.1
Expressions (TPMs) Base log 10
Testis
Embryo
Ovary
Brain
Gills
Heart
Muscle
Liver
Head kidney
Bones
Intestine
Danio rerio
Astyanax Mexicanus- cave
Anguilla anguilla
Astyanax Mexicanus- surface
Perca fluviatilis
Oncorhynchus mykiss

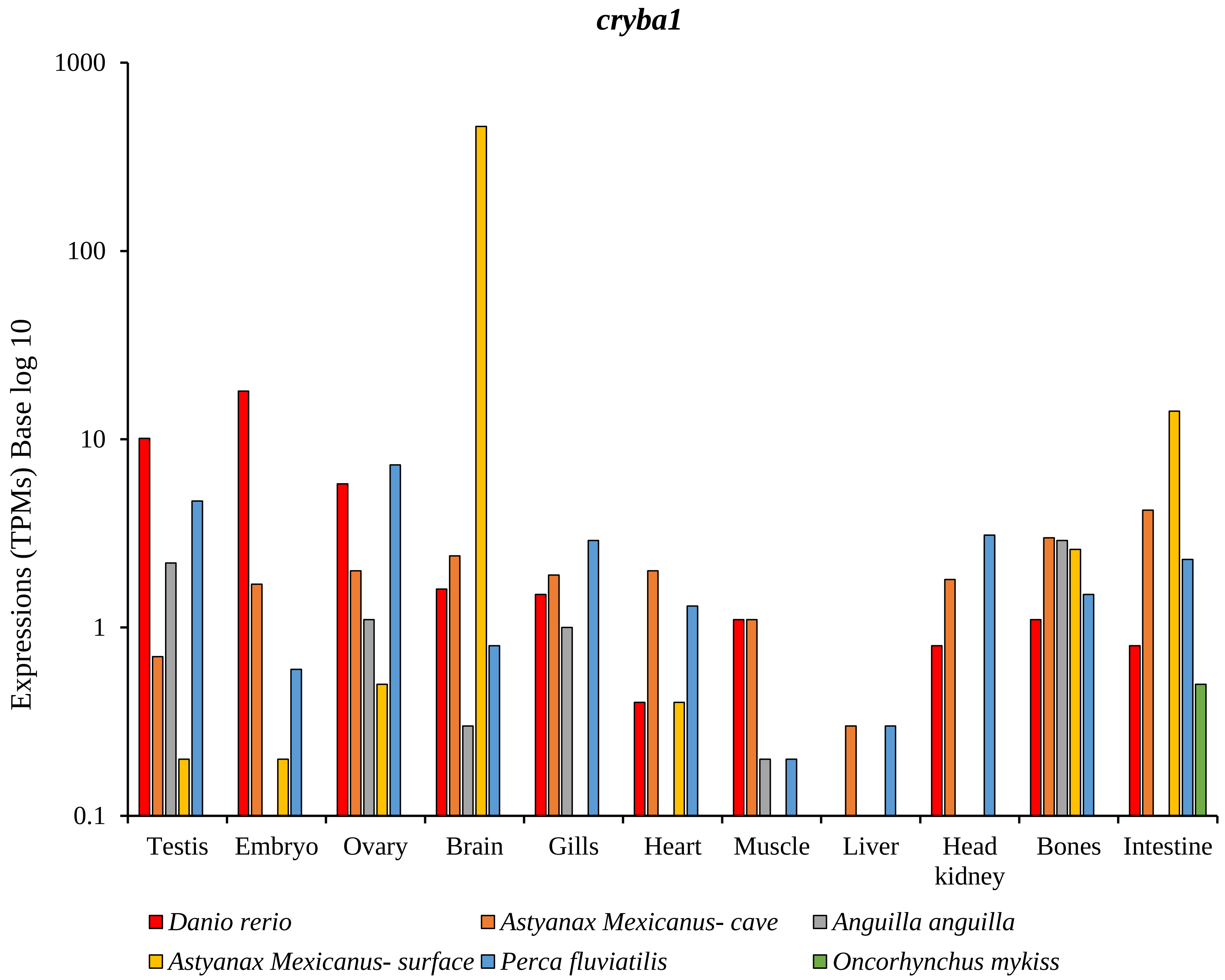

cryba1
Expressions (TPMs) Base log 10
1000
100
10
1
0.1
Testis
Embryo
Ovary
Brain
Gills
Heart
Muscle
Liver
Head kidney
Bones
Intestine
Danio rerio
Astyanax Mexicanus- cave
Anguilla anguilla
Astyanax Mexicanus- surface
Perca fluviatilis
Oncorhynchus mykiss

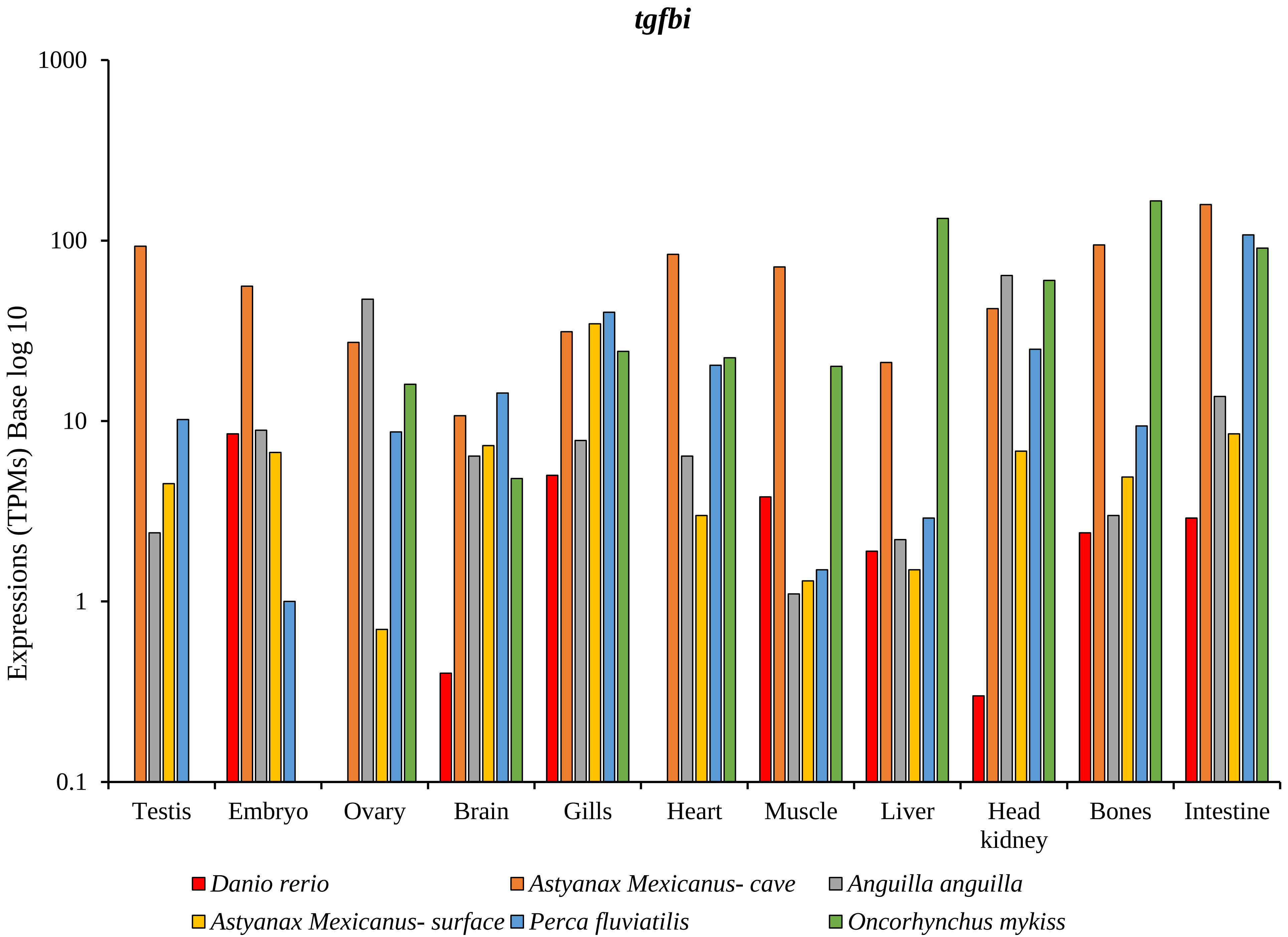

tgfbi
Expressions (TPMs) Base log 10
1000
100
10
1
0.1
Testis
Embryo
Ovary
Brain
Gills
Heart
Muscle
Liver
Head kidney
Bones
Intestine
Danio rerio
Astyanax Mexicanus- cave
Anguilla anguilla
Astyanax Mexicanus- surface
Perca fluviatilis
Oncorhynchus mykiss

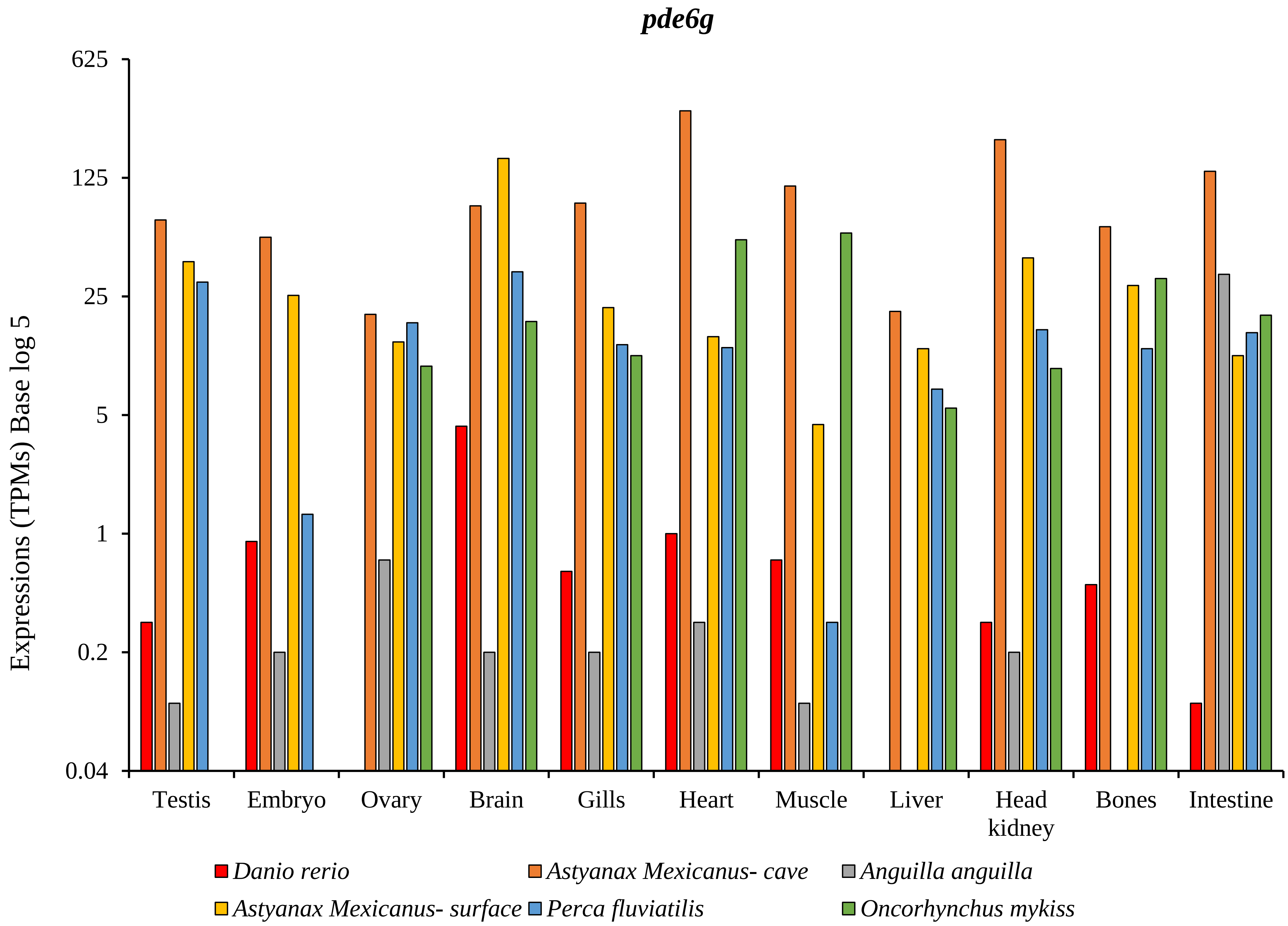
pde6g
Expressions (TPMs) Base log 5
625
125
25
5
1
0.2
0.04
Testis
Embryo
Ovary
Brain
Gills
Heart
Muscle
Liver
Head kidney
Bones
Intestine
Danio rerio
Astyanax Mexicanus- cave
Anguilla anguilla
Astyanax Mexicanus- surface
Perca fluviatilis
Oncorhynchus mykiss

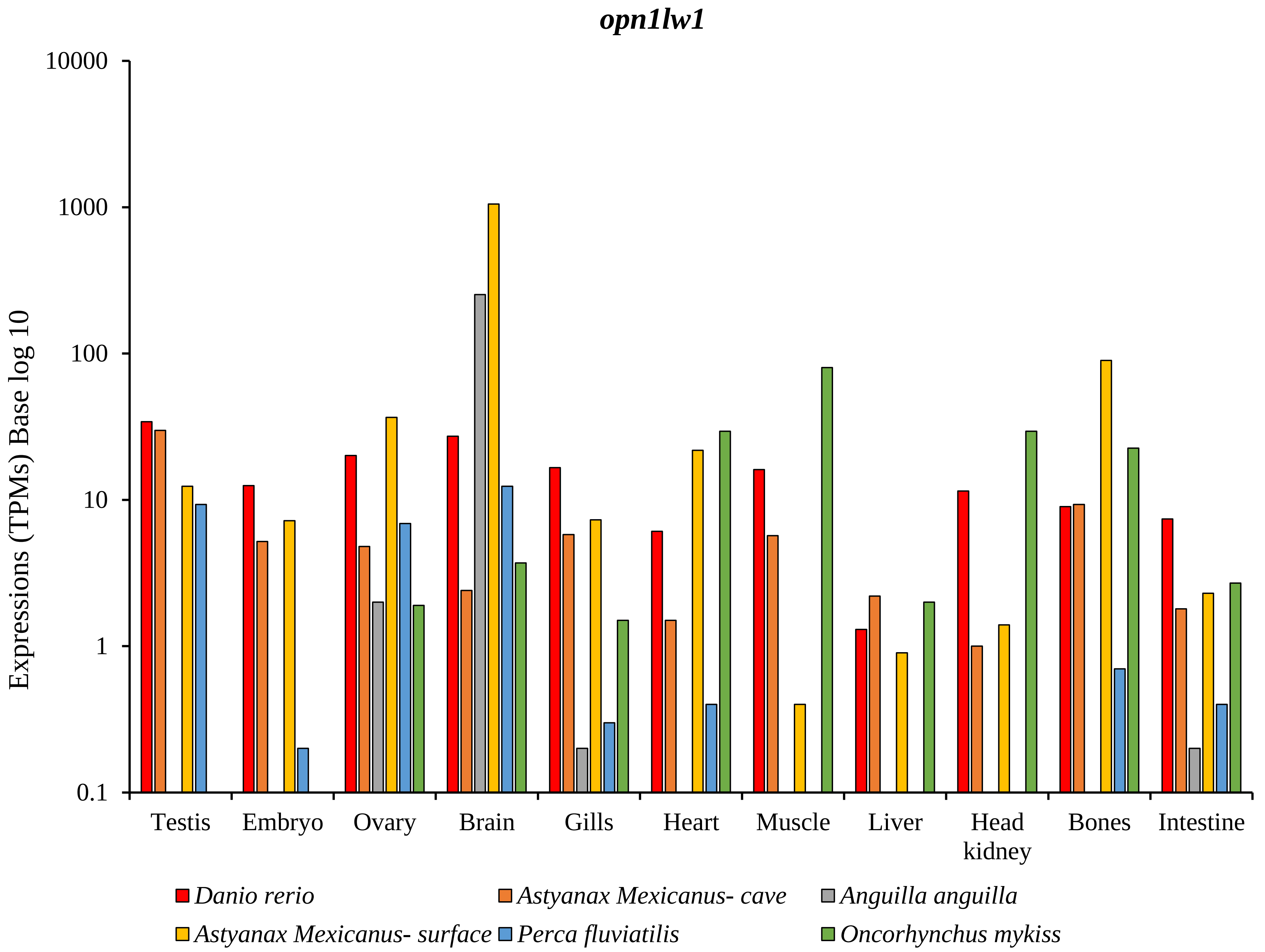
opn1lw1
Expressions (TPMs) Base log 10
10000
1000
100
10
1
0.1
Testis
Embryo
Ovary
Brain
Gills
Heart
Muscle
Liver
Head kidney
Bones
Intestine
Danio rerio
Astyanax Mexicanus- cave
Anguilla anguilla
Astyanax Mexicanus- surface
Perca fluviatilis
Oncorhynchus mykiss

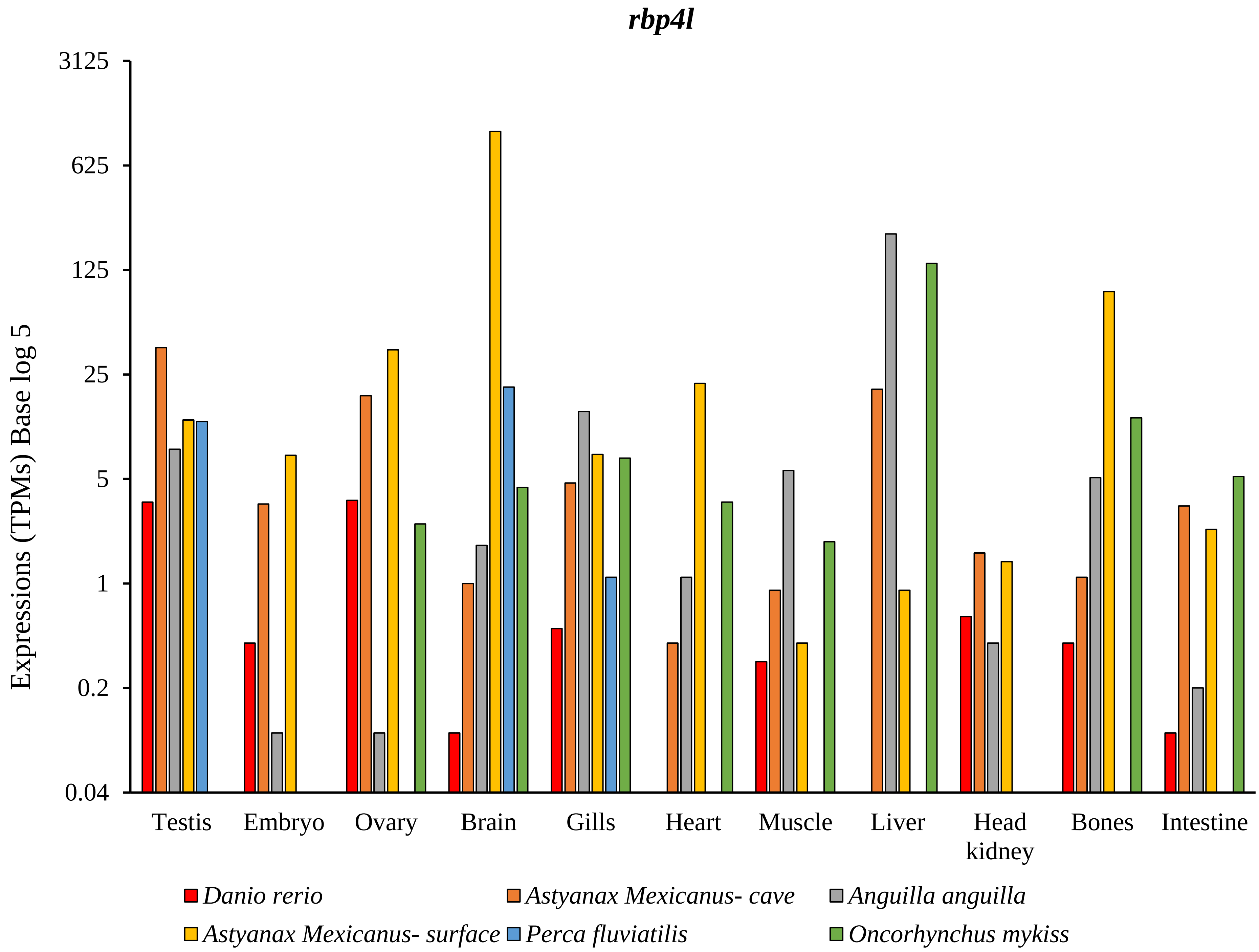
rbp4l
3125
625
125
25
5
1
0.2
0.04
Expressions (TPMs) Base log 5
Testis
Embryo
Ovary
Brain
Gills
Heart
Muscle
Liver
Head kidney
Bones
Intestine
Danio rerio
Astyanax Mexicanus- cave
Anguilla anguilla
Astyanax Mexicanus- surface
Perca fluviatilis
Oncorhynchus mykiss
