## Supplementary file 5 for "Paternal-effect genes revealed through semen cryopreservation in *Perca fluviatilis*"

| Gene name | Sequences | Amplicon length (bp) |
|---|---|---|
| crystallin, beta A2b [cryba2b] | F: ACAAGATCCGCTCCATCAAG | 167 |
| | R: GATGGGTCTGAAGGAAAGCA | |
| crystallin beta A4 [cryba4] | F: GCTATGAGCACGCCTCCTAC | 173 |
| | R: CTCACGCTCGTAGATGGTCA | |
| crystallin beta B1 [crybb1] | F: CATGATGTTCGACCAGGAGA | 153 |
| | R: TCCCCACGGAAGTTAGTCTG | |
| crystallin, gamma MX, like 2 [crygmxl2] | F: TAACTGCTGCAACTCCATGC | 134 |
| | R: AGTTGTTGAAGCCCATCCAG | |
| retinal cone rhodopsin-sensitive cGMP 3',5'-cyclic phosphodiesterase subunit gamma-like [pde6g] | F: AGACCGGACACAAACTGACC | 140 |
| | R: GGTCCTCTGCTTGAACTTGG | |
| red-sensitive opsin-like [opn1lw1] | F: CCAGGCGGTACAATGAAGAT | 105 |
| | R: GCGGAGCAATGTGGTAATTT | |
| gamma-crystallin M2-like [gamma M2] | F: GGGCAACCAGTACTTCCTGA | 188 |
| | R: CCATGACGTTGTCACAGTCC | |
| beta-crystallin A1-like [cryba1] | F: TACAGCGGTTCCCTCTCCTA | 218 |
| | R: AGCCAACTTCAGGCATCATC | |
| gamma-crystallin M3-like [crygm3] | F: GGAGAACTTCGGTGGTCAGA | 138 |
| | R: CCTCTGTAGTTGGGCTGCTC | |
| retinol binding protein 4, like [rbp4l] | F: TTTGACCCCAAGAGGTATGC | 165 |
| | R: ACACAACCCAGAAGCCAAAC | |
| transforming growth factor beta induced [tgfbi] | F: CTGAAGGAGCGTCTGTCCTC | 146 |
| | R: AAACGTCCGGTCTTATCGTG | |
| tetraspanin 7 [tspan7] | F: CACCAACTGCTCACCAGAGA | 179 |
| | R: ACAAGCAGCAGGACAGGAAT | |
| cytochrome c-like, transcript variant X1 [cycs] | F: TGTGGAGAATGGAGGAAAGC | 124 |
| | R: ATTCCAGACAATGCCTTTGC | |
| ER membrane protein complex subunit 10, transcript variant X2 [emc10] | F: GCCCAGCGTCTCACTAACTC | 172 |

| | | |
|---|---|---|
| | R: GGCTCTGACAAATGCTGTGA | |
| pre-mRNA-splicing factor [syf2] | F: GGAAGTTGTGGAGGAGGACA | 167 |
| | R: CTGCATCGTCAGCAGTGATT | |
| ER membrane protein complex subunit 3-like [emc3] | F: AACTGGGCCTTCTCTGGATT | 150 |
| | R: CCCAAACACGTTGAGGAAGT | |

**Table 1:** Details of the qRT PCR primers used in the study.
